# Transient YAP/TAZ inhibition exposes therapeutic vulnerabilities in advanced cancers

**DOI:** 10.1101/2025.07.03.662986

**Authors:** Marco Jessen, KyungMok Kim, Marie Tollot-Wegner, Anita Cindric Vranesic, Cagla Dönmez, Celina Junker, Tina Lehmann, Advitiya Khandelwal, Yuliya Kurlishchuk, Tom Hünniger, Christin Ritter, Evaristo Di Napoli, Shyam Krishnan Murali, Viktoria Buck, Sabine Muth, Tracy T. Tang, Andreas Rosenwald, Markus Radsak, Donato Inverso, Orlando Paciello, Björn von Eyss

**Affiliations:** Transcriptional Control of Tissue Homeostasis Lab, Leibniz Institute on Aging, Fritz Lipmann Institute e.V. (FLI); Jena, Germany; Institute of Pathology, University of Würzburg; Würzburg, Germany; Vascular Pathobiology Unit, Vita-Salute San Raffaele University; Milan, Italy; Department of Veterinary Medicine and Animal Production, University of Naples Federico II, Naples – Italy; Institute of Immunology and Research Center for Immunotherapy (FZI), University Medical Center of the Johannes Gutenberg-University Mainz; Mainz, Germany; Vivace Therapeutics, Inc.; San Mateo, USA; Department of Hematology and Oncology and Research Center for Immunotherapy (FZI), University Medical Center of the Johannes Gutenberg-University, Mainz, Germany; Department of Medicine VII - Hematology & Oncology, Donau-Isar-Klinikum, Deggendorf, Germany

## Abstract

YAP and TAZ, key effectors of the Hippo pathway, are frequently hyperactivated in cancer, where they drive tumor progression and resistance to therapy. Their oncogenic activity relies on interaction with TEAD transcription factors, making the TEAD-YAP/TAZ complex an attractive therapeutic target. Using translational mouse models, we demonstrate that sustained systemic YAP/TAZ depletion leads to severe side effects. However, even transient YAP/TAZ inhibition alone is sufficient to suppress tumor growth in advanced stages. Mechanistically, YAP/TAZ activity promotes T cell exclusion from the tumors by inducing target genes involved in tissue remodeling. Consequentially, YAP/TAZ inhibition induces immune infiltration, but the infiltrating T cells rapidly become exhausted. Combining YAP/TAZ inhibition with immune checkpoint blockade (ICB) overcomes this exhaustion and sensitizes previously resistant tumors to immunotherapy.

## Main text

### Ubiquitous long-term depletion of YAP/TAZ is highly detrimental

The Yes-associated protein (YAP) and transcriptional coactivator with PDZ-binding motif (TAZ/WWTR1) are the key signal transducers of the Hippo signaling pathway, which plays a critical role in regeneration (*1–3*). Dysregulation of YAP/TAZ activity, however, has been strongly implicated in cancer development, progression, and metastasis (*4, 5*). In numerous tumor types, YAP/TAZ are hyperactivated, promoting resistance to apoptosis, and a stem cell-like phenotype, contributing to aggressive tumor behavior and poor patient prognosis (*6*). Given their central role in oncogenic processes, YAP/TAZ have emerged as compelling therapeutic targets in oncology (*7, 8*). Since the oncogenic properties of YAP/TAZ - to a large extent - rely on their interaction with TEAD transcription factors (*8, 9*), many attempts are currently ongoing to design first compounds interfering with the transcriptional activity of the TEAD-YAP/TAZ complex as a novel target for tumor therapy (*10–13*). Given that the deletion of YAP/TAZ in adult mice under homeostatic conditions mostly does not result in significant adverse phenotypes (*14, 15*), the hypothesis that inhibitors targeting the TEAD-YAP/TAZ complex may have minimal side effects has been put forward.

In order to model the efficacy as well as potential adverse side effects of therapies targeting YAP/TAZ, we developed a doxycycline-inducible shRNA mouse model that enables the depletion of YAP/TAZ in the entire body of a mouse (Fig. 1A). An shRenilla (shRen) mouse line was used as non-targeting control as well as two shYAP/TAZ (shYT#1 and shYT#2) mouse lines, each expressing a distinct shRNA targeting *Yap* and *Taz (Wwtr1*), respectively. Both shYT lines showed a potent knockdown of YAP and TAZ, achieving approximately 80-90 % reduction across different tissues (Fig. 1B, fig. S1A-E). To validate the functionality of the shRNA mouse lines, we analyzed known YAP/TAZ-dependent knockout phenotypes in the shRNA lines: loss of biliary epithelial cells (BECs) and impairment of intestinal regeneration (Fig. 1C-G). Depletion of YAP/TAZ led to a significant reduction of BECs (Fig. 1C,D, fig. S2A,B) and strongly impaired intestinal regeneration after gamma irradiation (Fig. 1E-G). In all our experiments shYT#2 led to stronger phenotypes. These analyses thus demonstrate that the shRNA mouse model can recapitulate YAP/TAZ knockout phenotypes. Long-term depletion of YAP/TAZ (as compared to depletion of YAP or TAZ alone), was not well tolerated: most mice had to be sacrificed after one month due to excessive body weight loss (Fig. 1H, fig. S3A-G). Histological analysis at the humane endpoint revealed that shYT animals exhibited multifocal lesions across multiple organs, including hepatic inflammation, necrosis, and parenchymal degeneration; cardiomyocyte degeneration; and tubular epithelial cell damage in the kidney (Fig. 1I, fig. S4A-C). Thus, ubiquitous long-term YAP/TAZ depletion can lead to multi-organ defects. However, intermittent doxycycline administration could mitigate these adverse effects, enabling long-term experiments and suggesting the existence of a potential therapeutic window for YAP/TAZ inhibition (Fig. 1J).

**Fig. 1.**
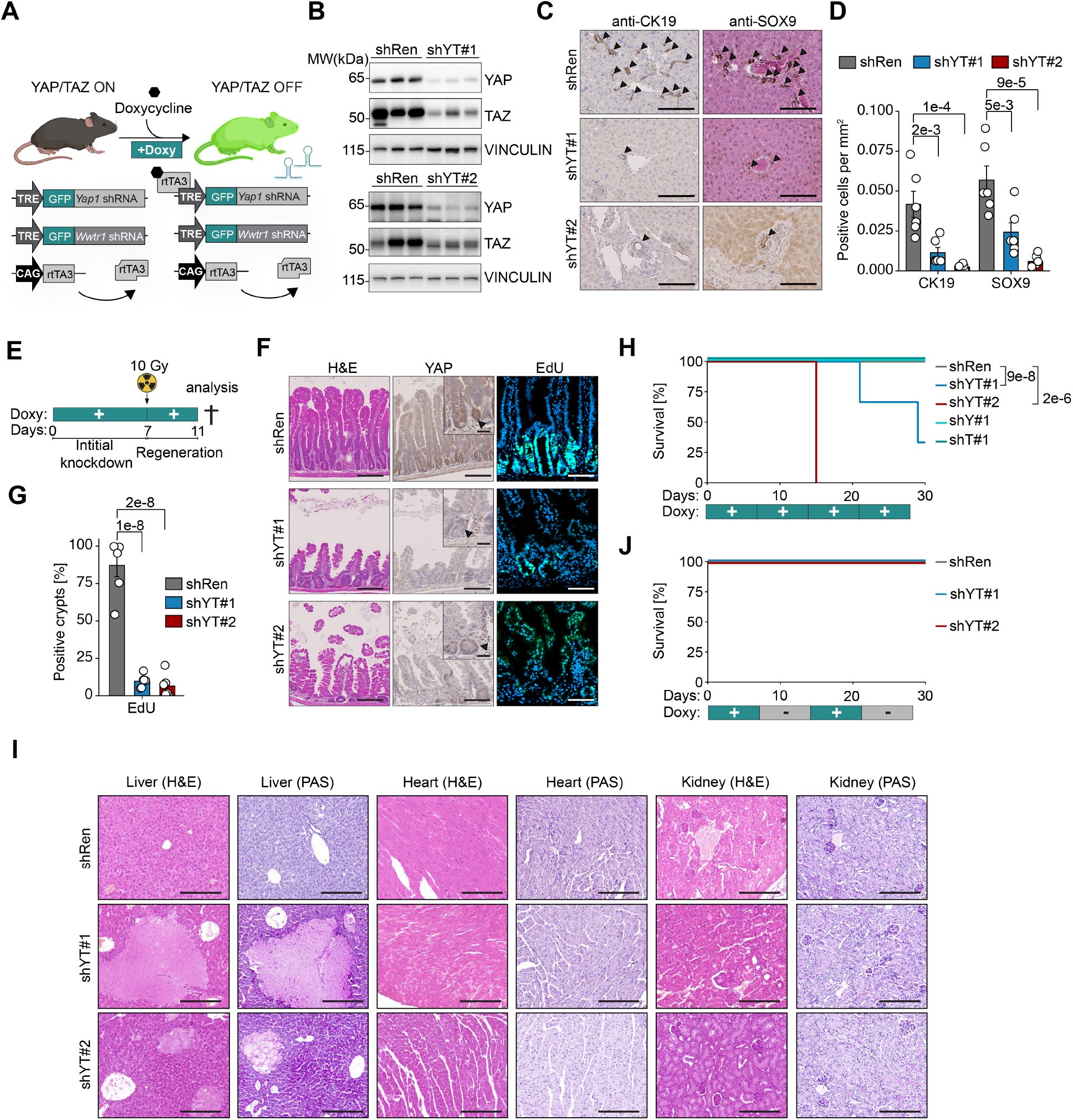
Systemic long-term depletion of YAP/TAZ is highly detrimental. **(A)** Schematic of the inducible shRNA mouse model. Ubiquitous expression of reverse tetracycline-controlled transactivator version 3 (rtTA3) under the CAG promoter enables ubiquitous expression of doxycycline-inducible shRNAs targeting *Yap1* and *Wwtr1* (Taz). Two independent mouse lines (shYT#1, shYT#2) were compared to control mice expressing shRNA against Renilla (shRen). **(B)** Western blot of liver lysates confirming YAP and TAZ knockdown after 2 weeks (shYT#2) or 4 weeks (shYT#1) of doxycycline treatment (n = 3 per group). (**C, D**) Liver sections stained for BEC markers CK19 and Sox9; arrows indicate counts of positive BECs. Quantification of BECs counts per liver area (n = 6 per group). Scale bars, 100 µm. **(E)** Experimental outline: after seven days of doxycycline-induced knockdown, mice were irradiated (10 Gy) and intestinal regeneration was assessed four days later. (**F, G**) Intestinal sections analyzed by H&E and YAP immunohistochemistry, and EdU (green) with DAPI (blue). Blowups highlight Paneth cells (arrows). Quantification of EdU-positive crypts (shRen, n = 6; shYT#1, n = 5; shYT#2, n = 6). Scale bars, 200 µm (overview), 50 µm (zoom). (**H, J**) Kaplan-Meier survival curves under constant (H) or intermittent (J) doxycycline administration for 4 weeks. Control group: shRen (n = 17 [H], n = 15 [J]); Additional mouse lines with shRNA targeting Yap1 (shY#1, n = 9) or Taz (shT#1, n = 11) alone are used as controls; shYT#1 (n = 6 [H], n = 7 [J]), shYT#2 (n = 6 [H], n = 4 [J]). **(I)** Representative hematoxylin and eosin (H&E) and periodic acid–Schiff (PAS) staining of liver, heart, and kidney sections from mice with the indicated genotypes after long-term depletion of YAP/TAZ. Scale bars, 200 µm. Data are means ± SEM. Statistical analyses: one-way ANOVA with Tukey HSD post hoc test (D, G) or ranked test (H).

### YAP/TAZ inhibition is beneficial in highly advanced cholangiocarcinoma

Cholangiocarcinomas (CCAs) are highly aggressive tumors with a poor prognosis for affected patients, and they show a critical dependency on high YAP/TAZ activity (*16*). We thus tested the applicability of ubiquitous YAP/TAZ inhibition in tumor therapy as a proof-of-concept in a CCA mouse model. Intermittent YAP/TAZ inhibition was sufficient to render mice tumor free in a hydrodynamic tail vein injection (HDTVI)-induced CCA model utilizing a transposase-based system to express activated versions of the Notch receptor (NICD) and myristoylated Akt (myr-AKT) (Fig. 2A-D). Next, we addressed how ubiquitous YAP/TAZ inhibition impacts CCA growth in a setting that more closely resembles a clinical setting: by administering only two weekly pulses of doxycycline in highly advanced tumors (Fig. 2E). This regimen was sufficient to nearly double the median survival of shYT animals (Fig 2F). The analysis of tumors from shYT animals - that reached the endpoint criteria after the second doxycycline pulse (“Late” timepoint) - showed a significant reduction of the liver to body weight ratio and tumor load demonstrating that transient YAP/TAZ inhibition is sufficient to severely reduce the tumor burden in highly advanced CCAs (Fig. 2F-I).

**Fig. 2.**
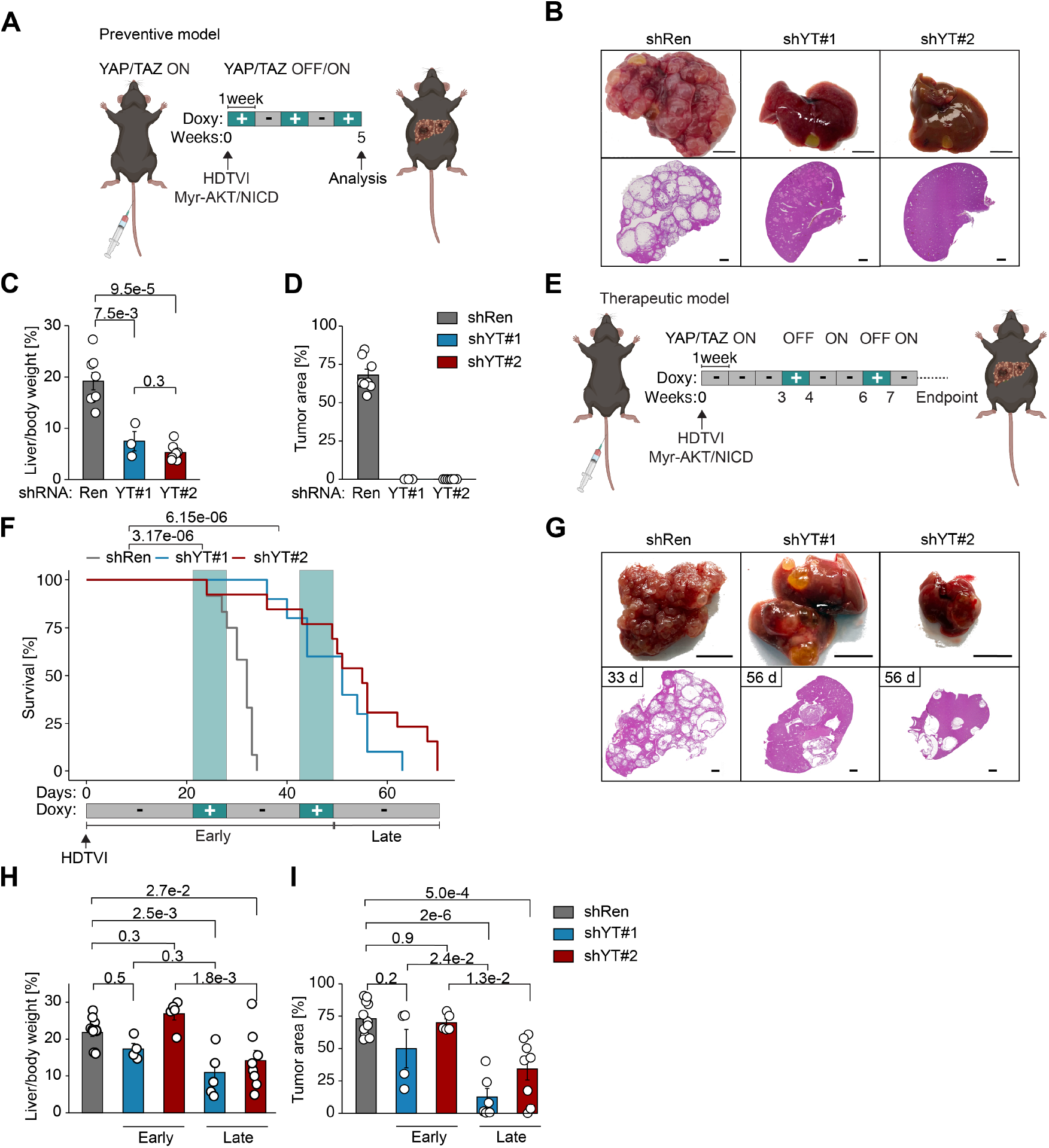
YAP/TAZ inhibition is beneficial in advanced cholangiocarcinoma. **(A)** Preventive model: YAP/TAZ depletion was initiated in the hydrodynamic tail vein injection (HDTVI) N-AKT cholangiocarcinoma model via weekly intermittent doxycycline administration. Tumors were analyzed five weeks after HDTVI. **(B)** Representative images (top) and H&E-stained liver sections (bottom) of indicated genotypes are shown. Scale bars, 1 cm (images), 2 mm (H&E). (**C, D**) Quantification of liver-to-body weight ratios and relative tumor area (shRen, n = 8; shYT#1, n = 3; shYT#2, n = 7). **(E)** Therapeutic model: YAP/TAZ depletion was transiently performed in advanced N-AKT cholangiocarcinoma at four and seven weeks after HDTVI. **(F)** Kaplan-Meier survival curves depict “Early” (week 1–7) and “Late” (week 8–10) survival of controls (shRen, n = 12) versus YAP/TAZ-depleted mice (shYT#1, n = 10; shYT#2, n = 13). **(G)** Representative livers (top) and H&E-stained liver sections are shown. The respective survival (in days) are indicated. Scale bars, 1 cm (images), 2 mm (H&E). (**H, I**) Quantification of liver-to-body weight ratios and relative tumor area (shRen, n = 12; shYT#1, n = 10; shYT#2, n = 13). Data represent means ± SEM. Statistical analyses: one-way ANOVA with Tukey HSD post hoc test (C, D, H, I) or ranked test (F).

### Acute YAP/TAZ depletion in advanced CCAs leads to T cell infiltration and activation

Next, we aimed to gain mechanistic insight into the early events triggered by YAP/TAZ inhibition in advanced tumors. To this end, we knocked down YAP/TAZ in advanced CCAs and analyzed the acute downstream effects (Fig. 3A). The mice were fed doxycycline starting from day 25 after HDTVI and the tumors were analyzed eight days later. Since it takes approximately five days for a knockdown to fully establish *in vivo* (*17*), the tumors have been depleted of YAP/TAZ for approximately three days. Consistently, YAP was barely detectable in shYT CCAs, *Yap1* and *Taz* mRNAs were strongly downregulated in our RNA-Seq data coinciding with downregulation of YAP/TAZ targets (Fig. 3B, fig. S5A, table S1). In the scRNA-Seq analyses (shRen: n=4, shYT#1: n=2, shYT#2: n=2), we could identify the tumor cells based on expression of AKT/NICD and annotate other cell types (Fig. 3C,D, fig. S6A). Using a highly conserved YAP/TAZ signature (*18*) to assess overall activity, we observed that tumor cells and cancer-associated fibroblasts (CAFs) exhibited the highest YAP/TAZ activity (Fig 3D). These cell types also showed the most pronounced changes in YAP/TAZ target gene expression following YAP/TAZ depletion (fig. S6B,C). Selective YAP/TAZ depletion in either CAFs or tumor cells (Fig. 3E), achieved via a Cre-inducible rtTA3 allele, revealed that targeting YAP/TAZ in tumor cells conferred a substantial survival advantage, whereas depletion in CAFs did not have a beneficial effect (Fig. 3F,G). T and NK cells which increased ~2-fold in shYT tumors, showed the most drastic relative increase of all cells within the tumor in shYT animals (Fig. 3H, fig. S7A). In addition, clusters of intratumoral T cells and CD8 cytotoxic T cells could only be observed in shYT tumors (Fig. 3I, J). T cells in shRen tumors could only be found sparsely at the tumor margin indicating that high YAP/TAZ plays an active role in T cell exclusion. Differential gene expression analyses within the T cell cluster revealed potent transcriptional responses indicative of T cell activation (Fig. 3K), arguing that YAP/TAZ depletion allows i) T cells to enter the tumor and ii) to become activated within in the tumor. An shYT T cell signature derived from the scRNA-Seq data set was highly predictive in a TCGA CCA data set, demonstrating that YAP/TAZ depletion can have a clinically important impact on the activities of intratumoral immune cells (Fig. 3L, table S2). An intercellular communication network analysis using CellChat (*19*) revealed that i) tumor cells showed enhanced interactions with T and NK cells and ii) T cells were predicted to increase their engagement with virtually all cell types (Fig. 3M, fig. S7B). Importantly, and in contrast to findings in other cancer types (*20*), YAP/TAZ knockdown did not alter Cd274 (PD-L1) expression in CCA tumor cells (fig. S7C).

**Fig. 3.**
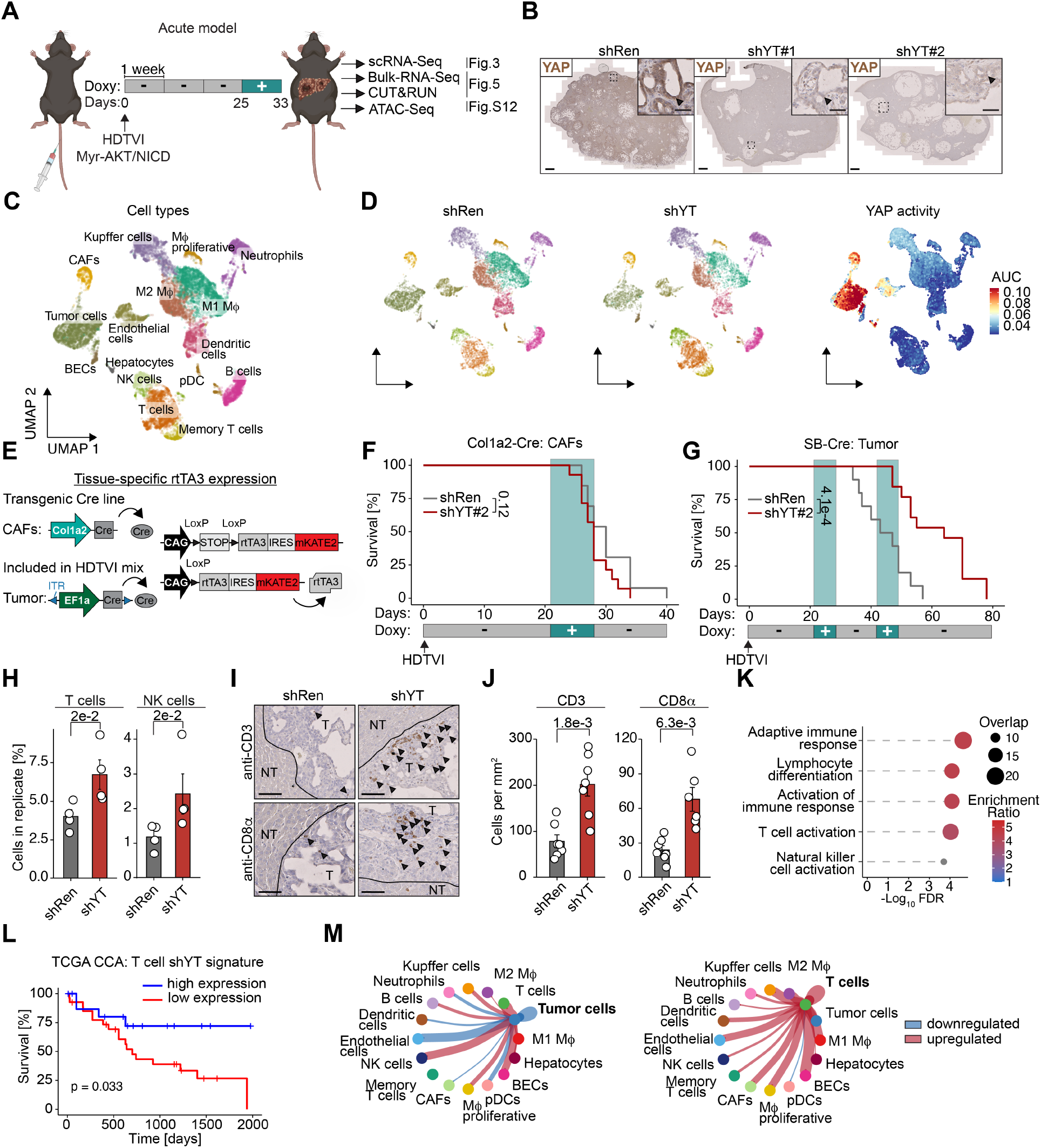
Acute YAP/TAZ depletion in advanced cholangiocarcinoma (CCA) promotes T cell infiltration and activation. **(A)** Acute depletion model: YAP/TAZ depletion was initiated at day 25 after HDTVI and analysed at day 33 in the N-AKT CCA model. **(B)** YAP immunohistochemistry (brown) in tumor-bearing livers. Arrows in the magnified view mark YAP-positive tumor cells. Scale bars: 1 mm (overviews), 50 µm (blowups). (**C, D**) UMAP plots from scRNA-seq following acute YAP/TAZ depletion: (C) overview of cell types in the N-AKT CCA model (D, left) Cell composition in shRen and shYT tumors; (D, right) YAP activation scores indicated across all cell types, based on a conserved YAP signature (*18*) (shRen, n = 4; shYT#1, n = 2; shYT#2, n = 2). (**E**) Schematic of tissue-specific rtTA3 expression (Rosa26-CAG-lsl-RIK). Cre-mediated deletion of the STOP cassette enables tissue-specific shRNA expression: in cancer-associated fibroblasts (CAFs; Col1a2-Cre) or tumor cells (Cre via plasmid in HDTVI). (**F, G**) Kaplan-Meier survival of mice with CAF-specific (shRen, n = 13; shYT#2, n = 14) or tumor cell-specific (shRen, n = 10; shYT#2, n = 13) YAP/TAZ knockdown. (**H**) Proportion of T cells (left) and NK cells (right), shown as percentage of total cells detected by scRNA-seq, following acute YAP/TAZ depletion (shRen, n = 4; shYT#1, n = 2; shYT#2, n = 2). (**I, J**) CD3 (top) and CD8a (bottom) immunostaining of CCAs after acute YAP/TAZ depletion, with quantification (shRen, n = 7; shYT#1, n = 2; shYT#2, n = 5). Arrows indicate positive cells. Scale bars, 100 µm. **(K)** Dot plot showing enriched pathways in YAP/TAZ-depleted T cells. **(L)** Kaplan-Meier survival of cholangiocarcinoma patients (TCGA dataset) stratified by high vs. low expression of a YAP/TAZ-dependent T cell activation signature. **(M)** Circle plots showing predicted ligand–receptor interactions of tumor cells (left) and T cells (right) with other cell types, inferred by CellChat. Line thickness indicates interaction strength; red and blue lines represent upregulated and downregulated interactions, respectively. Data represent means ± SEM. Statistical analyses: Welch‘s t test with Benjamini-Hochberg correction (H, J), log-rank test (F, G, L).

### YAP/TAZ depletion sensitizes CCAs towards immune checkpoint inhibitors

To test the contribution of CD8 cytotoxic T cells to the survival benefit in the therapeutic setting of shYT animals, we immunodepleted CD8 cytotoxic T cells during the two weekly doxycycline pulses (Fig 4A, fig. S8A,B). The depletion of CD8 cytotoxic T cells had no noticeable impact on the shRen control group indicative of a blunted immunosurveillance in the CCA model. In shYT#2 animals, however, the depletion of CD8 T cells significantly decreased the survival advantage indicating that YAP/TAZ depletion specifically activates CD8 cytotoxic T cells, contributing to the observed survival benefit (Fig. 4A,B).

**Fig. 4.**
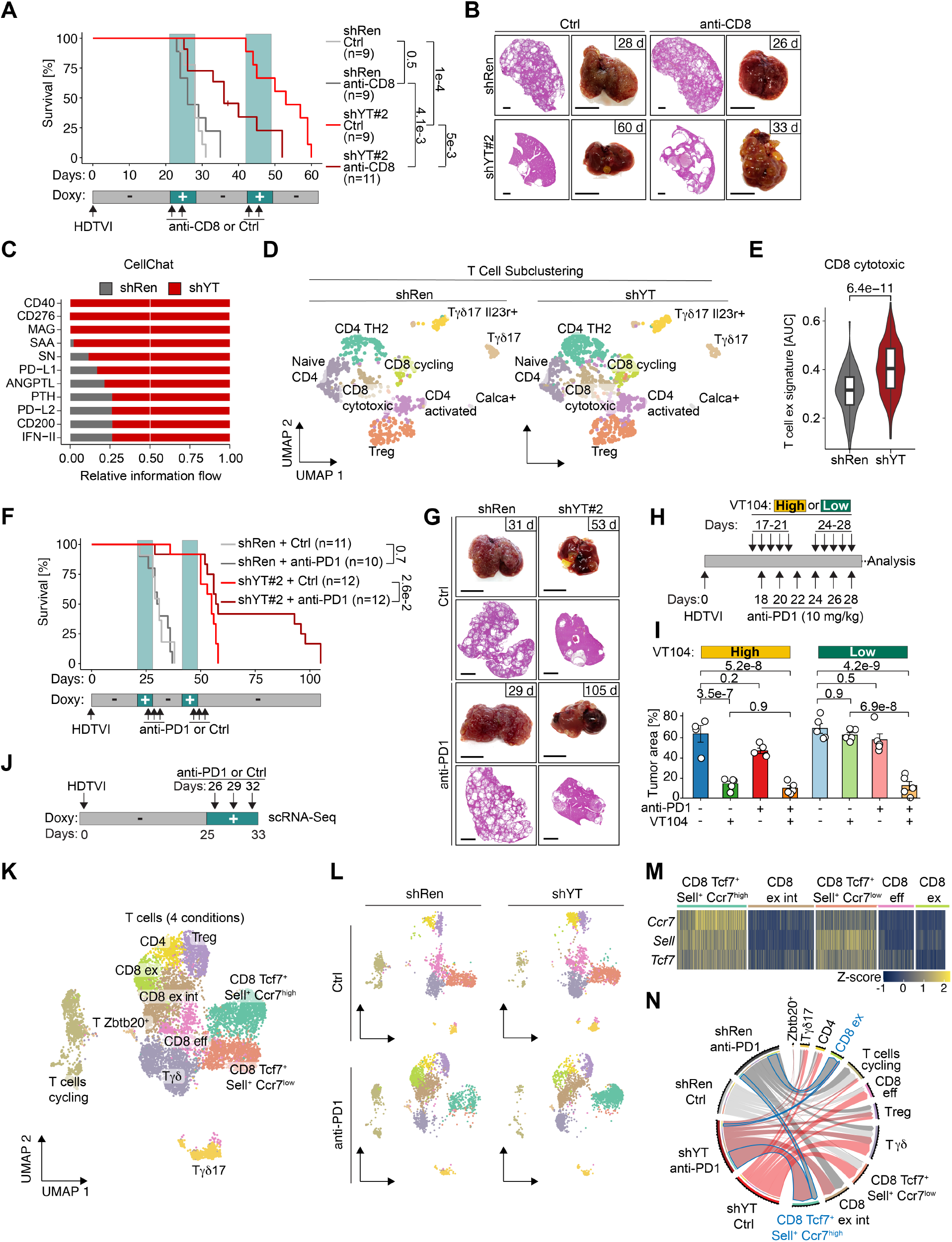
YAP/TAZ depletion sensitizes CCAs towards immune checkpoint blockade. (**A, B**) Kaplan-Meier survival curves of shRen and shYT#2 CCA-bearing mice with or without CD8 T cell depletion (shRen Ctrl, n = 9; shRen anti-CD8, n = 9; shYT#2 Ctrl, n = 9; shYT#2 anti-CD8, n = 11). Representative livers and H&E-stained liver sections are shown with the respective survival (in days). Scale bars, 1 cm (images), 1 mm (H&E). **(C)** CellChat analysis showing significantly upregulated ligand–receptor interactions upon YAP/TAZ depletion. X-axis indicates relative information flow, a quantitative measure of overall signaling activity. **(D)** UMAP plots illustrating reclustering of the T cell compartment from scRNA-seq data (Fig. 3). **(E)** Expression of a T cell exhaustion signature in CD8 cytotoxic T cells from shRen and shYT#2 tumors. (**F, G**) Kaplan-Meier survival curves of shRen and shYT#2 animals treated with or without anti–PD-1 (shRen Ctrl, n = 11; shRen anti–PD-1, n = 10; shYT#2 Ctrl, n = 12; shYT#2 anti–PD-1, n = 12). Representative livers and H&E-stained liver sections are shown with the respective survival (in days). Scale bars, 1 cm (gross images), 2 mm (H&E). (**H, I**) Schematic of a combination therapy with high (10 mg/kg) or low (1 mg/kg) doses of VT104 and anti–PD-1 in N-AKT–driven CCA. Tumor burden quantified in each group (n = 5; Ctrl (high-dose VT104), n = 4). (**J**) Schematic of acute YAP/TAZ depletion combined with anti–PD-1, followed by scRNA-seq analysis. (**K, L**) UMAP plots showing T cell clusters across all conditions (K) or stratified by treatment groups (L). **(M)** Heatmap for marker genes used for T cell subtype annotation of Tcf7^+^ Sell^+^ T cells. **(N)** Chord diagram showing relative abundance of T cell subtypes across treatment groups. The most prominent shifts in T cell populations are highlighted in blue. Data represent means ± SEM. Statistical analyses: Welch’s t test with Benjamini-Hochberg correction (E); one-way ANOVA with Tukey HSD post hoc test (I); log-rank test (A, F).

Further CellChat analyses revealed that the most significantly altered pathways in shYT animals were predominantly immune checkpoint-related - such as PD-L1, PD-L2, and CD276 - which became almost exclusively activated in this context (Fig. 4C). Subclustering of T cells corroborated that shYT tumors showed a strong increase in exhaustion signatures (Fig 4D,E, table S3). These findings suggest that although YAP/TAZ depletion triggers some level of T cell activation, immune checkpoints continue to restrain their full response, presenting an additional barrier to immune engagement. In the clinic, CCAs respond very poorly to immune checkpoint blockage (ICB), e.g. anti-PD1 antibodies which is largely attributed to the desmoplastic nature of CCAs as well as their immunosuppressive tumor microenvironment (*21*). From our previous analyses, we hypothesized that CD8 cytotoxic T cells infiltrating the tumor following YAP/TAZ depletion are rapidly suppressed and exhaust due to immune checkpoint activation. Based on this, ICB drugs should show a synergistic efficacy with YAP/TAZ depletion. To investigate this, we analyzed the survival of shRen and shYT#2 animals that were administered an anti-PD1 antibody targeting murine PD1. shRen animals did not show any measurable response to the anti-PD1 treatment whereas shYT#2 demonstrated a strong survival benefit with a high fraction of animals showing a doubled survival and reduced tumor load (Fig. 4F,G). These findings could be recapitulated with the TEAD inhibitor VT104 (*10*): at a high concentration (10 mg/kg), VT104 alone markedly reduced tumor burden, potentially reaching a therapeutic ceiling beyond which additional benefit from anti-PD1 treatment was not readily detectable. In contrast, at lower VT104 concentration (1 mg/kg), anti-PD1 treatment strongly synergized with VT104 to reduce tumor burden. (Fig. 4H,I, fig. S9A-C). To understand how YAP/TAZ depletion sensitizes tumors to ICB, we performed scRNA-Seq after YAP/TAZ depletion in combination with anti-PD1 (Fig. 4J). As expected, anti-PD1 led to a major shift in the T cell populations (Fig. 4K,L). Within the T cell population, we identified distinct subfractions of CD8 cytotoxic T cells exhibiting a progressive increase in exhaustion signatures (Fig. 4M, fig S10A-C, table S3, S4), transitioning from CD8 effector (eff) to CD8 exhausted intermediate (ex int), and ultimately to fully exhausted (ex) CD8 T cells. Tumors in shYT animals treated with anti-PD1 exhibited a marked increase in CD8 memory-like CD8+ Tcf7+ Sell+ Ccr7^high^ and a lower fraction of CD8 ex cells compared to shRen animals that also received anti-PD1 therapy (Fig. 4N). The fraction of intratumoral CD8 memory-like cells is a strong predictor for an efficient response to ICB (*22*) arguing that YAP/TAZ facilitates ICB response via this critical fraction of immune cells.

### YAP/TAZ drive a druggable oncogenic transcription program

Based on our findings, it is evident that TEAD-YAP/TAZ drive a transcriptional program which helps CCAs to grow aggressively, and to escape immunosurveillance, e.g. by driving T cell exclusion. By comparing transcriptional profiles of tumor cells with their non-malignant counterparts - BECs and hepatocytes - we found that YAP/TAZ activate a tumor-specific transcriptional program absent in non-malignant cells (fig. S11A, table S5). To identify the primary drivers of this oncogenic TEAD-YAP/TAZ program, we performed CUT&RUN analysis for TEAD1 and YAP in EpCAM-enriched tumor cells acutely depleted of YAP/TAZ (fig. S11B-C). These data were integrated with bulk RNA-seq and ATAC-seq datasets generated from the same purified cells (Fig. 5A). TEAD1 CUT&RUN experiments revealed 3,390 high-confidence peaks (q-value < 1e–4), strongly enriched for TEA domain motifs (fig. S11B). Since TEAD proteins preferentially bind to enhancer regions (Fig. 5B), directly linking peaks to gene regulation is challenging. To address this, we used ATAC-Seq to assess chromatin accessibility and validate the functional relevance of TEAD1 binding sites. Notably, TEAD motifs were highly enriched in regions that became less accessible following YAP/TAZ depletion (fig. S12A–C). Based on the cumulative distribution frequency (CDF) of TEAD1 peaks, we inferred that genes located within 10^7^ bp of a TEAD1 peak likely represent the majority of direct targets (fig. S11C). In the bulk RNA-Seq dataset, gene ontology (GO) terms associated with tissue remodeling were significantly enriched among the downregulated genes, which was accompanied by a marked reduction in intratumoral collagen and loss of the desmoplastic nature (Fig. 5C-E, fig. S13A, B). Since this suggested that YAP/TAZ regulate a critical set of direct targets involved in tissue remodeling - ultimately contributing to T cell exclusion we compiled a list of 144 direct TEAD1 targets that are responsive to YAP/TAZ depletion (fig. S11D, table S6). Further filtering using GO terms for “enzymatic activity” and “extracellular” identified 70 potentially druggable candidates from which we selected eight known to be involved in T cell exclusion (Fig. 5F). These genes were knocked out after HDTVI *in vivo*, using a doxycyclin-inducible Cas9 mouse line (iCas9) (Fig. 5G). The most prominent hits were *Tgfb2* and *Enpp3*, as their knockout was as effective as targeting the primary oncogenic driver myr-AKT (Fig. 5H, I, fig. S14,15). This analysis uncovers a set of pivotal target genes that function as central mediators of YAP/TAZ-driven tissue remodeling and oncogenesis *in vivo*.

**Fig. 5.**
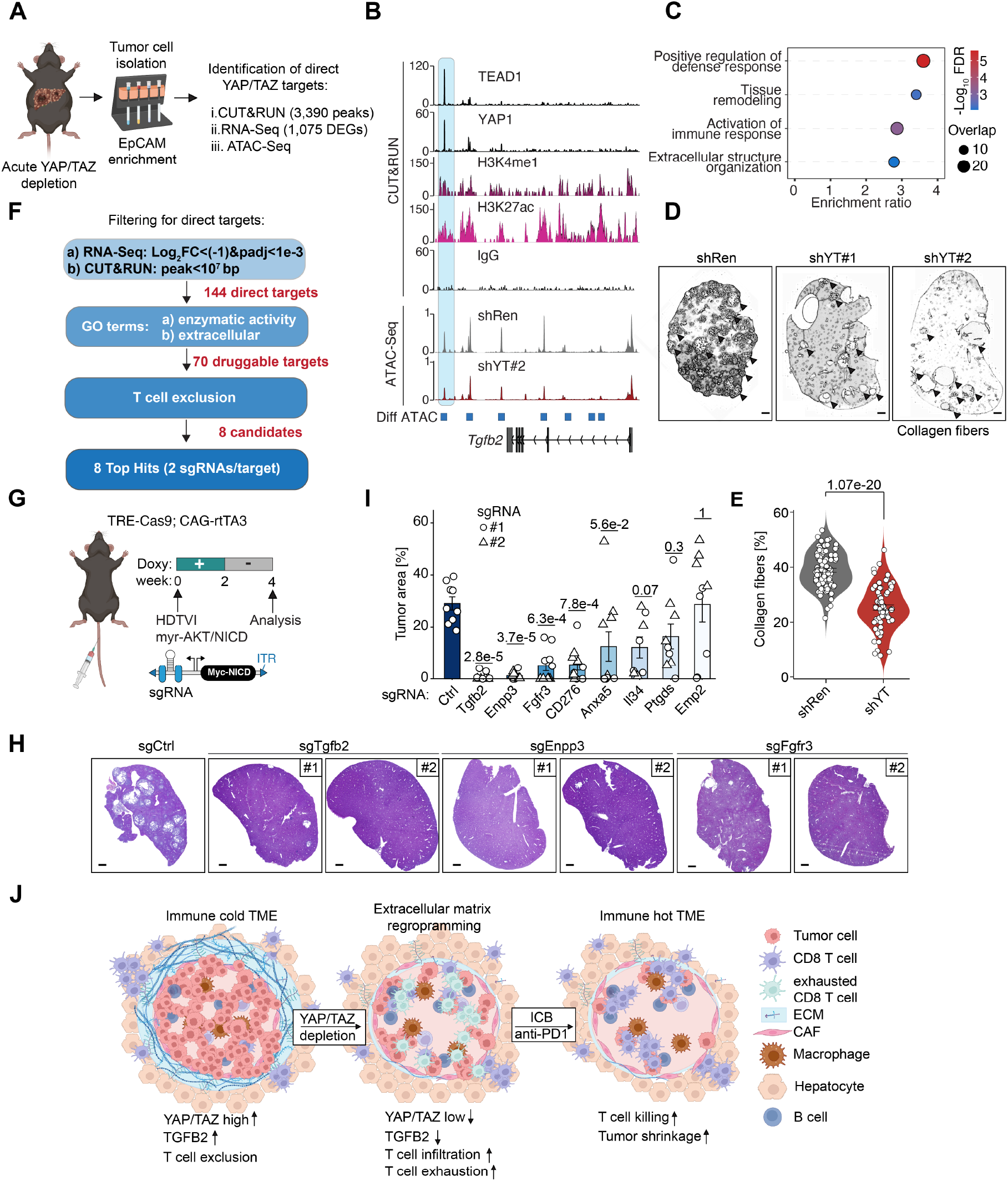
YAP/TAZ drive a druggable oncogenic transcriptional program. **(A)** Schematic outlining the identification of direct YAP/TAZ target genes in cholangiocarcinoma cells. **(B)** Genome browser tracks at the *Tgfb2* locus showing TEAD1, YAP1, H3K4me1, H3K27Ac, and IgG CUT&RUN profiles, along with ATAC-seq data, in control (shRen) and YAP/TAZ-depleted (shYT#2) tumor cells. Diff ATAC = Differential ATAC-Seq peaks that are down is shYT tumor cells. **(C)** Dot plot for a pathway enrichment analysis, highlighting immune activation and extracellular matrix organization pathways upregulated upon acute YAP/TAZ depletion. (**D, E**) Representative WEKA-segmented images of Sirius Red–stained liver sections, highlighting collagen fibers (black) and background tissue (white) in tumors with acute YAP/TAZ depletion (shYT#1, n = 2; shYT#2, n = 5) or control (shRen, n = 6). Collagen content was quantified. Scale bars, 100 µm. (**G**) Schematic of an inducible Cas9 mouse model combined with hydrodynamic tail vein injection (HDTVI) of sgRNA-expressing plasmids to evaluate the functional role of candidate YAP/TAZ targets in N-AKT–driven CCA growth. (**H, I**) Representative H&E-stained liver sections of the indicated gene knockouts with quantification of tumor area. Scale bars, 1 mm. (n = 5, per group; n = 4, sgIl34#1, #2; n = 9, sgCtrl) (**J**) Two-hit model for the synergy between YAP/TAZ depletion and ICB: YAP/TAZ depletion remodels the extracellular matrix, facilitating T cell infiltration and subsequent exhaustion. ICB reverses this exhaustion, rendering the CCA tumor microenvironment (TME) responsive to ICB. Data represent means ± SEM. Statistical analyses: One-way ANOVA with Tukey HSD post hoc test (E, I).

Our findings demonstrate that ubiquitous long-term YAP/TAZ inhibition causes severe side effects, e.g. by impacting vital organs like the heart and liver. Notably, pulsed inhibition preserved anti-tumor efficacy while minimizing toxicity. This suggests that, although not without consequence, YAP/TAZ inhibition has a broader therapeutic window
- especially when combined with immune checkpoint blockade (ICB). The synergy between YAP/TAZ depletion and ICB can be explained by a two-hit model: YAP/TAZ reversed by ICB (Fig. 5J). Thus, YAP/TAZ inhibitors may render ICB-resistant tumors responsive to immunotherapy.

## Acknowledgements

The DNA sequencing, the proteomics, the imaging, the flow cytometry, and the mouse facilities as well as the core service histology of the FLI are gratefully acknowledged. We would like to thank all the members of the von Eyss lab, von Maltzahn lab, and Kaether lab for helpful discussion. We thank Tom Hünniger and Christin Ritter and the entire animal care taker team at FLI - especially Jenny Buchelt, Claudia Maisch, Björn Brüns, and Patrick Elsner - for their excellent technical support. We thank the entire animal welfare team — especially Patricia Schmitt — for their valuable support and guidance regarding animal licences. We would like to thank Tae-Won Kang (Zender lab) for hydrodynamic tail vein injection training. Some of the figures were created with BioRender.com.

## Funding

Wilhelm Sander-Stiftung 2022.084.1(BvE) BMBF 16GW0271K (BvE)

DFG EY 120/4-1 (BvE) DFG EY 120/9-1 (BvE) DFG EY 120/10-1 (BvE)

German Cancer Aid/Deutsche Krebshilfe 70116078 (BvE) Leibniz Collaborative Excellence K398/2021

DFG Project-ID 318346496 - SFB1292 subproject TP21N (SM and MPR)

## Author contributions

Conceptualization: MJ, BvE Methodology: MJ, BvE, SM, MR, TT

Investigation: MJ, VB, DI, OP, AR, KMK, MTW, ACV, CD, CJ, TL, AK, YK, TH, CR, EDN, SKM

Visualization: MJ, BvE Funding acquisition: BvE,

Project administration: BvE Supervision: BvE

Writing – original draft: MJ, BvE Writing – review & editing: MJ, be

