## Supplementary figures and methods for "Transient YAP/TAZ inhibition exposes therapeutic vulnerabilities in advanced cancers"

**The PDF file includes:**

Materials and Methods
Figs. S1 to S15
Tables S7 to S13

**Other Supplementary Materials for this manuscript include the following:**

Tables S1 to S6

### Materials and Methods

#### Mouse Strains

The TRE-tGFP-shYAP1.351 (shY#1), TRE-tGFP-shYAP1.322 (shY#2), TRE-tGFP-shWwtr1.2056 (shT#1), TRE-tGFP-shWwtr1.4587 (shT#2), and TRE-tGFP-shRen mouse lines were generated and purchased from Mirimus. In these lines, tGFP serves as a fluorescent marker, and a specific shRNA is expressed from a promoter containing a tetracycline response element (TRE). The knock-in allele is located downstream of the *Col1a1* locus (17). The ubiquitous doxycycline-inducible expression of the shRNAs is facilitated by the reverse tetracycline-controlled transactivator 3 (rtTA3), which is driven by a CAG promoter. The animals used in the experiments carry knock-in alleles that express shRNAs targeting *Yap1* and *Taz*, while the control animals carry knock-in alleles expressing an shRNA targeting Renilla luciferase. To ensure tissue-specific expression of the shRNAs, a Rosa26-lox-Stop-lox (LSL) cassette upstream of rtTA3 was used (23). CAF-specific expression was achieved by crossing the shRNA lines with Col1a2-Cre transgenic mice (24). For tumor cell-specific shRNA expression, Cre recombinase delivered through the hydrodynamic tail vein injection (HDTVI) plasmid mix. iCas9 animals were purchased from JAX (#029476). These animals harbor an inducible TRE-Cas9 allele which is downstream of the *Col1a1* locus. When combined with the CAG-rtTA3 transgene, doxycycline administration enables ubiquitous expression of Cas9. Control animals were sex- and age-matched. Mice were housed, fed, and treated according to the guidelines of the Leibniz Institute on Aging - Fritz-Lipmann Institute (FLI). All animal experiments were approved by the federal state of Thuringia and conducted in accordance with national guidelines and regulations. Animal experiments presented in this manuscript were performed under the following approved licenses: FLI-17-017, FLI-21-015, FLI-22-012, FLI-23-017, and FLI-24-002.

Experimental scoring procedure

Animals undergoing long-term YAP/TAZ depletion studies were weighed and scored weekly if no clinical score was applied. Animals exhibiting a score of 1 or 2 were monitored daily, while a score of 3 was defined as the humane endpoint, at which the experiment was terminated.

The predefined scoring system was as follows:

- Score 0: body weight unchanged
- Score 1: 0–10% loss of body weight
- Score 2: 10–20% loss of body weight
- Score 3: Over 20% loss of body weight

Animals reaching the humane endpoint criteria were defined as those with more than 20% body weight loss compared to their baseline weight at the start of the experiment.

Animals undergoing cholangiocarcinoma (CCA) tumor studies were scored weekly from days 0 to 25, and then every two to three days from day 25 onward, unless a clinical score was applied. Animals exhibiting a score of 1 or higher were monitored daily, while a score of 3 was defined as the humane endpoint, at which point the experiment was terminated.

The predefined scoring system was as follows:

- Score 0: Normal body condition score
- Score 1: Mild abdominal protrusion/swelling over the liver region
- Score 3: Pronounced abdominal distension and enlargement of the abdomen without fluid accumulation (ascites), with palpable individual liver tumors

In addition to the specific scoring criteria for this study, animals were also scored according to the animal welfare guidelines of the Leibniz Institute on Aging - Fritz Lipmann Institute (FLI). This included assessments of general condition and grooming, body condition score, posture and signs of pain, behavior, eyes and body openings, as well as fur and skin condition.

### Plasmids

The pCMV(CAT)T7-SB100 plasmid that expressed hyperactivate sleeping beauty transposase was purchased from addgene (#34879). Myristolated, HA-tagged Akt, Myc-tagged Notch1 intracellular domain (NICD) were amplified by PCR from pT3-myr-AKT-HA (Addgene #31789), pT3-EF1aH NICD (Addgene #86500) and cloned into the SfiI restriction site of the pSBbi-Pur vector (Addgene #60523). The puromycin resistance was removed from the sleeping beauty vectors by EcoRI/BsrGI restriction digest followed by blunting and ligation.

To generate a bispecific Sleeping Beauty vector that expresses Myc-tagged NICD under the control of the EF1α promoter and a U6-driven sgRNA cassette, the following fragments were cloned by Gibson assembly into the SfiI/EcoRI restriction sites of the pSBbi-Pur vector (Addgene #60523): the EF1α promoter was amplified by PCR from the pSBbi-Pur vector (Addgene #60523), Myc-tagged NICD was amplified from pT3-EF1aH NICD (Addgene #86500), and the U6-SapI-sgRNA cassette was amplified from PX552 (Addgene #60958).

### sgRNA design

The ChopChop tool was used for sgRNA design (25). sgRNAs were cloned into the pSBbi-Myc-NICD-sgRNA vector (cut with SapI) and Gibson assembly of a 70-mer oligo following the design: ATCTTGTGGAAAGGACGAAACACCG, 20 nt sgRNA, GTTTTAGAGCTAGAAATAGCAAGTT.

### Generation of mouse embryonic fibroblasts (MEFs)

Mouse embryonic fibroblasts (MEFs) were isolated from pregnant mice at embryonic day E13.5. After removing the liver and head, embryos were minced and digested with 0.25% trypsin. Following 15 minutes of incubation, the embryos were pipetted up and down vigorously, and fresh trypsin was added. After additional 15 minutes, the digested embryos were plated in DMEM medium containing 10% fetal bovine serum and 1%

Penicillin/Streptomycin. For immortalization, MEFs were infected with a retrovirus containing an shRNA targeting p19 ARF.

#### Hydrodynamic tail vein injection

For the delivery of Sleeping Beauty-based plasmids to the murine liver, hydrodynamic tail vein injection was performed. Mice were anesthetized using isoflurane. Specifically, 25 µg of pSBbi-Myr-AKT-HA, 25 µg of pSBbi-Myc-NICD, and 10 µg of pCMV-SB100 plasmids were diluted in sterile Ringer-Lactate solution. The total volume was adjusted to 10% of the animal's body weight and injected within 6-8 seconds. Mice were kept on a heating plate at 37°C until they recovered from anesthesia. Mice were monitored for 3 hours after HDTVI and then daily until they reached appropriate time points.

#### Doxycycline administration

To induce expression of shRNAs in shYT animals or Cas9 in iCas9 animals, mice were fed doxycycline-containing chow (625 mg/kg doxycycline hyclate; Sniff, #326652) for the indicated duration.

#### Anti-CD8, anti-PD1 administration

To deplete $CD8^+$ T cells, animals were injected intraperitoneally with 500 µg of a neutralizing monoclonal anti-CD8 antibody (clone YTS169) at the indicated timepoints. Successful $CD8^+$ T cell depletion was confirmed by flow cytometric analysis of peripheral blood. An anti-mouse PD-1 antibody (clone RMP1-14, BioXcell, #BE0146) was administered intraperitoneally at a dose of 10 mg/kg body weight at the indicated timepoints.

#### Peripheral blood analysis

To confirm efficient depletion of $CD8^+$ T cells, 25 µL of peripheral whole blood was incubated for 30 minutes with fluorochrome-conjugated antibodies (1:100 dilution in FACS buffer) targeting CD11b (myeloid marker), CD3 (T cell marker), B220 (B cell marker), CD8α (CD8 T cells), and CD4 (CD4 T cells) (Table S7). Following antibody incubation, red blood cells were lysed, and cells were washed twice with FACS buffer. Dead cells were excluded by staining with Sytox Blue. Samples were analyzed on a Cytek Spectral Flow Cytometer.

#### VT104 administration

VT104 (Vivace Therapeutics) was freshly formulated daily as a suspension in a vehicle composed of 5% DMSO, 10% Solutol (Sigma Aldrich), and 85% glucose (5%) in water (D5W). To prepare the formulation, 3.35 mg of VT104 powder was weighed into a vial and mixed with 50 µL DMSO by vortexing for 5 minutes. Subsequently, 100 µL Solutol was added and vortexed for an additional 2 minutes. The suspension was sonicated for

10 minutes, followed by the addition of 850 µL D5W and vortexed for 2 minutes. A final 10-minute sonication ensured a homogeneous suspension of approximately 1 mL (~3.3 mg/mL). To achieve the target concentration of 2 mg/mL, 667 µL of vehicle was added, resulting in a total volume of ~1.67 mL—sufficient to dose five mice at 10 mg/kg with a dosing volume of 5 µL/g body weight. The formulation was prepared immediately prior to administration and used within 1–2 hours at room temperature. For lower dose administrations (1 mg/kg), the 10 mg/kg stock solution was diluted 1:10 with vehicle and administered accordingly. VT104 was administered by oral gavage within 2 hours of formulation.

#### Whole body irradiation and EdU administration

To assess the effect of YAP/TAZ depletion on small intestine regeneration, animals were exposed to a sublethal dose of gamma irradiation (10 Gy) using a GC40 irradiation chamber (Caesium-137 source). After 4 days of regeneration, animals were sacrificed, and intestinal regeneration was evaluated. To quantify proliferating cells, animals were injected intraperitoneally with EdU (10 mM stock solution; 100 µL per 10 g body weight) 2 hours prior to sacrifice.

#### Histology, immunohistochemistry and image analysis

Fixation, paraffin embedding and H&E staining

Mouse livers were fixed in 10% formalin (3.7% formaldehyde in 1× PBS) for at least 48 hours at 4 °C. Following fixation, livers were dehydrated and embedded in paraffin. Paraffin sections (4 µm thickness) were deparaffinized and processed for either hematoxylin and eosin (H&E) staining or immunohistochemistry (IHC) using specific primary antibodies.

Immunohistochemistry

For IHC, antigen retrieval was performed using either citrate buffer (pH 6) or Tris-EDTA (TE) buffer (pH 9), depending on the antibody. Sections were boiled in a Coplin jar using a microwave for 12 minutes, followed by a 30-minute incubation at room temperature. After cooling down, sections were blocked in 3% bovine serum albumin (BSA) in PBST (PBS with 0.1% Tween-20) for 2 hours at room temperature. Primary antibodies were diluted in blocking buffer to the appropriate concentrations and incubated on the sections overnight at 4 °C. The following primary antibodies were used: YAP1 (Cell Signaling #14074; citrate pH 6; 1:200), Sox9 (Millipore #5535; citrate pH 6; 1:200), CK19 (DSHB TROMA-III; TE pH 9; 1:15), CD3 (Abcam ab16669; citrate pH 6; 1:150), CD8 (Cell Signaling #98941S; citrate pH 6; 1:200). The following day, sections were washed three times for 5 minutes each in PBST and incubated with a secondary antibody conjugated to horseradish peroxidase (HRP; 1:1000 in PBST) for 1 hour at room temperature. After washing, visualization was performed using DAB chromogen (Vector Labs) according to the manufacturer's protocol, followed by counterstaining with hematoxylin. Sections were then dehydrated and mounted using a xylene-based mounting medium.

EdU staining

EdU staining was performed using the Click-iT™ EdU Cell Proliferation Kit (Thermo Fisher) according to the manufacturer's instructions. Slides were imaged using an automated Axio Scan.Z1 microscope (Zeiss).

Picro-Sirius Red Staining for Collagen Visualization

To visualize collagen fibers in cholangiocarcinoma (CCA) tumor sections, Picro-Sirius Red staining was performed. Tissue sections were first deparaffinized as previously described. Nuclear counterstaining was carried out using Weigert's hematoxylin for 8 minutes, followed by a 10-minute rinse under running tap water. Collagen fibers were stained with a saturated Picro-Sirius Red solution (0.5 g Sirius Red dissolved in 500 mL saturated aqueous picric acid) for 1 hour at room temperature. Sections were then rinsed twice with acidified water (prepared by adding 5 mL glacial acetic acid to 1 L of tap water) to remove excess dye. Finally, sections were dehydrated through graded alcohols, cleared in xylene, and mounted with a xylene-based mounting medium.

Periodic acid-Schiff (PAS) staining

The Periodic acid–Schiff (PAS) reaction was used to detect polysaccharides, e.g. glycogen. Tissue sections were deparaffinized, rehydrated through a graded ethanol series, and incubated in 0.5% periodic acid solution for 5–10 minutes to oxidize glycols to aldehydes. Sections were then rinsed in distilled water and treated with Schiff's reagent for 15–20 minutes, followed by thorough washing in running tap water to develop the magenta color. Nuclei were counterstained with hematoxylin. Then, sections were evaluated, scanned, and photographed using the Pannoramic II scanner (3DHISTECH, The Digital Pathology Company, Övutca 3, 1141 Budapest, Hungary).

Histological scoring of liver, kidney and heart

A morphological evaluation was performed using haematoxylin and eosin staining to identify tissue alterations across different organs. In the kidneys, the focus was on detecting signs of tubular cell degeneration. Liver sections were examined for the presence of inflammation, hepatocyte degeneration, and necrosis. In the heart, attention was given to identifying degenerative changes and necrosis in cardiomyocytes. The lesions (tubular cell degeneration, hepatic inflammation, hepatocyte degeneration, hepatic necrosis, degenerative changes, and necrosis in cardiomyocytes) were evaluated by examining at least 10 microscopic fields at 20× magnification, using histological scoring systems. In particular, score 0 (absent lesions or no inflammatory foci); score 1 (mild lesions or <2 foci per 20× field); score 2 (moderate lesions or 2–4 foci per 20× field); and score 3 (severe lesions or >4 foci per 20× field.

**Cholangiocarcinoma cell (CCA) isolation**

Tumor-bearing livers were minced and digested in 15 mL of 1× PBS containing 7.5 mg Pronase, 7.5 mg Collagenase P, and 1.5 mg DNase I using a C Tube (Miltenyi Biotec) and the gentleMACS Octo Dissociator (Miltenyi) for 45 minutes at 100 rpm and 37 °C. Following digestion, the cell suspension was filtered through a 100 µm cell strainer and washed with an appropriate volume of Advanced DMEM/F12 (AdDMEM/F12) medium. Red blood cell lysis was performed using the Red Blood Cell Lysis Solution (Miltenyi,

#130-094-183), followed by debris removal (Miltenyi, #130-109-398) and dead cell removal (Miltenyi, #130-090-101) according to the manufacturer's protocols. Cells were then incubated with Fc receptor blocking reagent (Miltenyi) for 10 minutes on ice. Tumor cells were labeled with anti-EpCAM-APC antibody (eBioscience, #17-5791-82; 1:100) for 30 minutes on ice. After washing, cells were incubated with anti-APC microbeads (Miltenyi) for an additional 30 minutes. The cell suspension was then loaded onto LS columns (Miltenyi) and processed according to the manufacturer's instructions. $EpCAM^+$ cells were eluted and further processed for downstream applications.

#### Biliary epithelial cell (BEC) isolation

Liver tissues were finely minced and processed following the procedure described under the Cholangiocarcinoma Cell Isolation section. Red blood cells were removed using red blood cell lysis buffer (Miltenyi). To enrich for non-parenchymal cells, hepatocytes were depleted via low-speed centrifugation (5 minutes at 30 ×g). The remaining cell suspension was incubated with Fc receptor blocking reagent (Miltenyi Biotec) for 10 minutes on ice to prevent nonspecific antibody binding. BECs were identified and labeled using an anti-EpCAM-APC antibody (eBioscience, #17-5791-82; dilution 1:100). To exclude hematopoietic and erythroid lineage cells, samples were simultaneously stained with the following lineage markers (LIN) conjugated to APC-Cy7: TER-119 (erythroid cells; BD Biosciences, #560509, 1:100), CD11b (myeloid cells; BD Biosciences, #557657, 1:100), CD45 (pan-leukocyte marker; BD Biosciences, #557659, 1:100). Staining was performed for 30 minutes on ice in the dark. After two washes with FACS buffer (PBS + 2% FBS), samples were filtered through a 40 µm mesh and analyzed on a BD cell sorter. BECs were defined as $EpCAM^+/LIN^-$ events.

#### RNA isolation and Quantitative RT-PCR

For RNA isolation from cultured cells, cells were lysed in TRIzol reagent (VWR). For tissue samples, the corresponding tissue was minced, resuspended in TRIzol, and mechanically homogenized using the Precellys Tissue Homogenizer (Bertin Technologies). Following lysis, samples were processed according to the manufacturer's protocol. Briefly, RNA was separated from DNA and proteins by phenol-chloroform extraction and subsequently precipitated using isopropanol. The concentration and purity of RNA were assessed using a NanoDrop 1000 spectrophotometer (Thermo Fisher Scientific), and samples were then processed for downstream applications.

For quantitative real-time PCR (RT-qPCR), 2 µg of total RNA was reverse transcribed into cDNA using M-MLV Reverse Transcriptase (Promega) following the manufacturer's instructions. The resulting cDNA was diluted 1:15, and RT-qPCR was performed using the InnuMIX qPCR MasterMix (Analytik Jena) in accordance with the manufacturer's specifications.

### Whole protein lysate and immunoblots

For whole-cell lysates, cells or tissues were lysed directly in ice-cold RIPA buffer (50 mM Tris-HCl pH 7.4, 150 mM NaCl, 1% NP-40, 0.5% sodium deoxycholate, 0.1% SDS) supplemented with a protease inhibitor cocktail (Sigma-Aldrich) for 10 minutes on ice. For tissue samples, minced tissue was homogenized using the Precellys Tissue Homogenizer (Bertin Technologies) in RIPA buffer. Lysates were centrifuged at 16,000 × g for 10 minutes at 4°C, and the supernatant containing soluble proteins was collected.

Protein concentrations were determined using the BCA Protein Assay Kit. Samples were mixed with 5× SDS loading buffer and denatured for 5 minutes at 95°C. Equal amounts of protein (10–25 µg) were separated on 8% Bis-Tris gels for 2 hours at 120 V and transferred onto 0.45 µm PVDF membranes (Millipore) for 90 minutes at 250 mA.

Membranes were blocked in 5% non-fat dry milk prepared in TBS containing 0.25% Tween-20 (TBS-T) for at least 30 minutes at room temperature with constant agitation. Primary antibody incubation was performed overnight at 4°C in 5% BSA in TBS-T using 50 mL conical tubes under continuous rotation. The following day, membranes were washed three times with TBS-T and incubated with horseradish peroxidase (HRP)-conjugated secondary antibodies for 1 hour at room temperature. After three additional washes with TBS-T, chemiluminescent signals were developed using Clarity ECL substrate (Bio-Rad) and imaged.

### Bulk RNA-Seq analysis

For bulk RNA sequencing, TRIZOL-purified total RNA was additionally purified using the RNeasy Mini Kit (Qiagen) including an on-column DNase I digestion for 15 minutes at room temperature to remove residual genomic DNA. RNA quality and concentration were assessed using the Fragment Analyzer (Agilent Technologies). For library preparation, 1 µg of high-quality RNA was used as input for mRNA enrichment using the NEBNext® Poly(A) mRNA Magnetic Isolation Module (New England Biolabs). mRNA was then processed using the NEBNext Ultra II RNA Library Prep Kit in combination with NEBNext Multiplex Oligos for Illumina (E6440, New England Biolabs), following the manufacturer's protocol. Final libraries were quantified using the Agilent 2100 Bioanalyzer (Agilent Technologies) and sequenced as 75 bp single-end reads on an Illumina NextSeq 500 platform. Sequencing reads were extracted in FASTQ format using bcl2fastq (v1.8.4, Illumina). Adapter trimming, size selection (retaining reads >25 nucleotides), and quality filtering (Phred score >30) of FASTQ files were performed using Cutadapt. Reads were subsequently aligned to the mouse genome (mm10) with Bowtie2 (version 2.2.9) using default settings. Read counts were extracted in R using the countOverlaps function from the GenomicRanges package. Differential gene expression analysis was carried out with DESeq2 (version 3.26.8) using default parametersv.

**scRNA-Seq analysis**

Effect of acute YAP/TAZ-depletion on TME (10x Genomics)

For single-cell RNA sequencing, tumor-bearing livers were digested and processed as described in the Cholangiocarcinoma Cell (CCA) Isolation protocol. Following dead cell removal, viable cells were incubated with Fc receptor blocking reagent (Miltenyi Biotec) for 10 minutes on ice to prevent non-specific antibody binding. Subsequently, cells were labeled with 1 µg of anti-mouse hashtag antibodies (TotalSeq™-B, BioLegend) for 30 minutes on ice. After labeling, cells were washed twice with PBS and resuspended in 1× PBS. The final cell concentration was adjusted to 1,000 cells/µL for downstream single-cell library preparation. Single-cell encapsulation and barcoding were performed using the 10x Genomics Chromium Controller in combination with the Chromium Next GEM Single Cell 3' Reagent Kits v3.1 (10x Genomics), according to the manufacturer's protocol. Libraries were sequenced on an Illumina platform with a targeted depth of ~50,000 reads per cell. Raw sequencing data were processed using Cell Ranger (v6.1.2) (10x Genomics) with the mm10 mouse reference genome for demultiplexing, alignment, and generation of gene-cell count matrices. Downstream analysis was performed using the Seurat R package (v4.3). Cells with fewer than 500 detected genes, more than 5,000 detected genes, or with >10% mitochondrial gene expression were excluded from further analysis. Hashtag oligonucleotide (HTO)-based sample demultiplexing was performed using Seurat's HTODemux function. After quality control, datasets were normalized using SCTransform, and samples were integrated using Seurat's integration workflow based on canonical correlation analysis (CCA). Dimensionality reduction was performed using principal component analysis (PCA), followed by UMAP for visualization. Clustering was conducted using a shared nearest neighbor (SNN) graph and the Louvain algorithm.

Effect of acute YAP/TAZ-depletion and anti-PD1 treatment on TME (Singleron)

Cell isolation and processing were performed as previously described in the 10x Genomics section, with the exception that hashtag antibody labeling was omitted. Following cell isolation, the final cell concentration was adjusted to 750 cells/µL for downstream single-cell library preparation. Single-cell encapsulation, barcoding, and cDNA synthesis were performed using the Singleron GEXSCOPE Single Cell RNA Library Kit V2, according to the manufacturer's instructions. Prepared libraries were sequenced on an Illumina platform, targeting a read depth of approximately 25,000 reads per cell. Raw sequencing data were processed using Cell Scope (Singleron) with the mm10 (mouse) reference genome, which included steps for sample demultiplexing, read alignment, and generation of gene-cell count matrices. Downstream analysis (e.g., quality control, clustering, and differential expression analysis) was conducted as described previously in the 10x Genomics section.

**CUT&RUN**

Cholangiocarcinoma (CCA) cells were isolated as previously described (see Cholangiocarcinoma Cell Isolation). Following EpCAM enrichment, tumor cells were counted and resuspended at a concentration of 2 × $10^6$ cells in wash buffer (20 mM HEPES, pH 7.5; 150 mM NaCl; 0.5 mM spermidine; protease inhibitor cocktail). For each

CUT&RUN reaction, 2 × $10^5$ cells were used. Cells were incubated with 10 µL of activated Concanavalin A-coated magnetic beads (Polysciences) for 10 minutes at room temperature on a hula mixer with constant agitation. Following bead binding, samples were placed on a magnetic stand to remove the supernatant. Bead-bound cells were resuspended in 100 µL of antibody buffer (20 mM HEPES, pH 7.5; 150 mM NaCl; 0.5 mM spermidine; 0.01% [w/v] digitonin; 2 mM EDTA; protease inhibitor), and primary antibodies were added as follows: anti-TEAD1 (Cell Signaling Technology, #12292), anti-YAP1 (Cell Signaling Technology, #14074), anti-H3K27Ac (Abcam, ab4729), and anti-H3K4me1 (Cell Signaling Technology, #5326). A species-matched IgG was used as a negative control. Samples were incubated overnight at 4°C on a HulaMixer (Thermo Fisher Scientific) set at a 50° tilt and 1 RPM. The next day, samples were washed twice with digitonin wash buffer (20 mM HEPES, pH 7.5; 150 mM NaCl; 0.5 mM spermidine; 0.01% digitonin; protease inhibitor). Protein A/G-MNase (pAG-MNase; 1 µg/mL) was added in digitonin wash buffer, and samples were incubated for 1 hour at 4°C with gentle agitation. After incubation, cells were washed three times with digitonin wash buffer and twice with low-salt buffer (20 mM HEPES, pH 7.5; 0.5 mM spermidine; 0.01% digitonin). Nuclease activation was performed by transferring the samples to calcium-containing buffer (3.5 mM HEPES, pH 7.5; 10 mM $CaCl_2$; 0.01% digitonin; protease inhibitor) and incubating on ice for 1 hour with continuous mixing. Cleaved DNA fragments were released by incubation at 37°C for 30 minutes in STOP buffer (170 mM NaCl; 20 mM EGTA; 0.01% digitonin; 50 µg/mL RNase A; 25 µg/mL Glycogen Blue). Supernatants were collected, and DNA was purified by phenol-chloroform extraction. Library preparation was performed using the NEBNext Ultra II DNA Library Prep Kit for Illumina (NEB, #E7645) following the manufacturer's protocol. Library quality was assessed using the Fragment Analyzer (Agilent Technologies), and equimolar pooling of libraries was performed. Pooled libraries were gel purified, quantified on an Agilent 2100 Bioanalyzer, and sequenced as 75 bp paired-end reads on an Illumina NextSeq 500 platform.
CUT&RUN analysis was conducted as previously described (26). Briefly, adapter and quality trimming were performed using Cutadapt. Reads were aligned to the mm10 genome and a repeat-masked *E. coli* genome (used as spike-in) using Bowtie2. Reads with inserts <120 bp mapped to mm10 were extracted (deepTools), and a scaling factor based on the *E. coli* spike-in was used to generate normalized bigWig files. Peak calling was performed using Genrich using the following parameters: q 1e-4 -a 700. For cumulative distribution frequency (CDF) analysis, TEAD1 signals at categorized peaks (based on RNA-Seq regulation, downregulated, upregulated or not-regulated) were associated with the nearest TSS using BEDtools, and plotted with ggecdf (ggpubr).

### OMNI ATAC-Seq

Cholangiocarcinoma (CCA) cells were isolated as described (see Cholangiocarcinoma Cell Isolation). A total of 5 × $10^4$ cells were pelleted in a pre-wetted 1.5 mL low-binding tube by centrifugation at 500 × g for 5 minutes at 4°C using a fixed-angle rotor. Cells were resuspended in 50 µL ice-cold ATAC-Resuspension Buffer (RSB: 10 mM Tris-HCl, pH 7.4; 10 mM NaCl; 3 mM $MgCl_2$) supplemented with 0.1% (v/v) NP-40, 0.1% (v/v) Tween-20, and 0.01% (w/v) digitonin. The suspension was pipetted up and down three times and incubated on ice for 3 minutes to lyse the cells. To remove the lysis reagents, 1 mL ice-cold RSB containing 0.1% Tween-20 (without NP-40 or digitonin) was added, and the tube was gently inverted three times. Nuclei were pelleted by centrifugation at 500 × g for 10 minutes at 4°C, and the supernatant was carefully removed. Nuclei were resuspended in 50 µL of transposition reaction mix containing: 25 µL 2× TD buffer (Illumina), 2.5 µL Tn5 transposase (to reach 100 nM final concentration), 16.5 µL 1× PBS, 0.5 µL 1% digitonin, 0.5 µL 10% Tween-20, 5 µL nuclease-free water. The mix was pipetted up and down six times to ensure homogeneous suspension. Samples were incubated at 37°C for 30 minutes at 1000 rpm using a thermomixer. Following transposition, DNA was purified using the Zymo DNA Clean & Concentrator-5 Kit (Zymo Research) according to the manufacturer's instructions. Libraries were amplified using the NEBNext 2× Master Mix (New England Biolabs) and assessed for quality using the Fragment Analyzer (Agilent Technologies). Final libraries were pooled in equimolar ratios and sequenced as single-end reads on the Illumina NextSeq 500 platform. For the ATAC-Seq analysis the nextflow pipeline (https://nf-co.re/atacseq/2.1.2/) was used. The differential peak analysis was performed using DiffChIPL (https://github.com/yancychy/DiffChIPL) and TF motif accessibility was analyzed with TOBIAS (27).

#### Gene set enrichment analysis (GSEA)

Gene set enrichment was performed using the MSigDB GSEA tool (v 4.3.1) with a GSEAPreranked analysis.

#### Quantifications and statistics

Statistical analyses were conducted using R (version 4.1.0). Unless otherwise specified, graphs depict the mean along with the standard error of the mean (SEM). The specific statistical test applied is indicated in the corresponding figure legend. Each experiment was performed with a minimum of three independent biological replicates, unless stated otherwise in the figure legend.

Image quantifications were performed in a blinded manner. Tumor area, collagen content, and the desmoplastic index were assessed using H&E-stained sections (for tumor area and desmoplastic index) or Picro Sirius Red–stained sections (for collagen fibers). Images were analyzed using ImageJ in combination with the Trainable Weka Segmentation plugin. The area of all tumor nodules and collagen fibers was measured in pixels and normalized to the total liver area. To calculate the desmoplastic index, the percentage of tumor stroma was measured in pixels, and normalized by the total tumor area.

**Data and materials availability:** The Next-generation sequencing data generated in this study have been deposited in the GEO database under accession code GSE301501. The histology images have been deposited in the BioImage Archive under accession code S-BIAD2127.

### Supplemental Figures

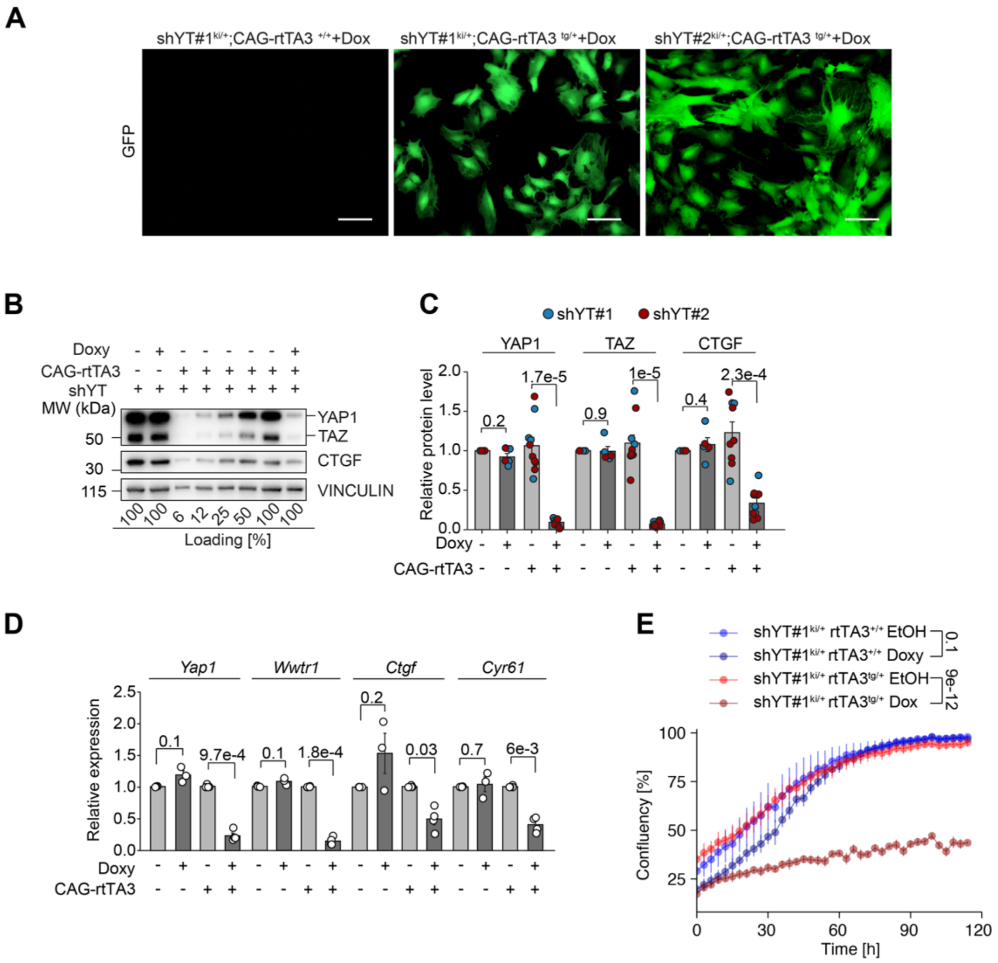


**Fig. S1. Validation of YAP/TAZ knockdown in mouse embryonic fibroblasts (MEFs).** (**A**) Representative images of MEFs shYT#1, shYT#2 mouse lines or control MEFs of the same lines lacking the reverse tetracycline transactivator (rtTA3), showing GFP expression. Scale bars, 50 µm.
(**B**, **C**) Representative immunoblots of MEF lysates harvested after 5 days of doxycycline-induced shRNA expression, showing protein levels of YAP1, TAZ, and CTGF. Control lysates were serially diluted to better quantify knockdown efficiency. Quantification across replicates (n = 3–6 per condition).
(**D**) RT–qPCR analysis of *Yap1*, *Wwtr1*, *Ctgf*, and *Cyr61* expression in shYT#1 MEFs.
(**E**) Proliferation of shYT#1 MEF lines assessed by confluency measurements using an Incucyte imaging system.
Data represent mean ± SEM. Statistical analysis: Welch's t test with Benjamini-Hochberg correction (C, D); one-way ANOVA with Tukey HSD post hoc test (E).

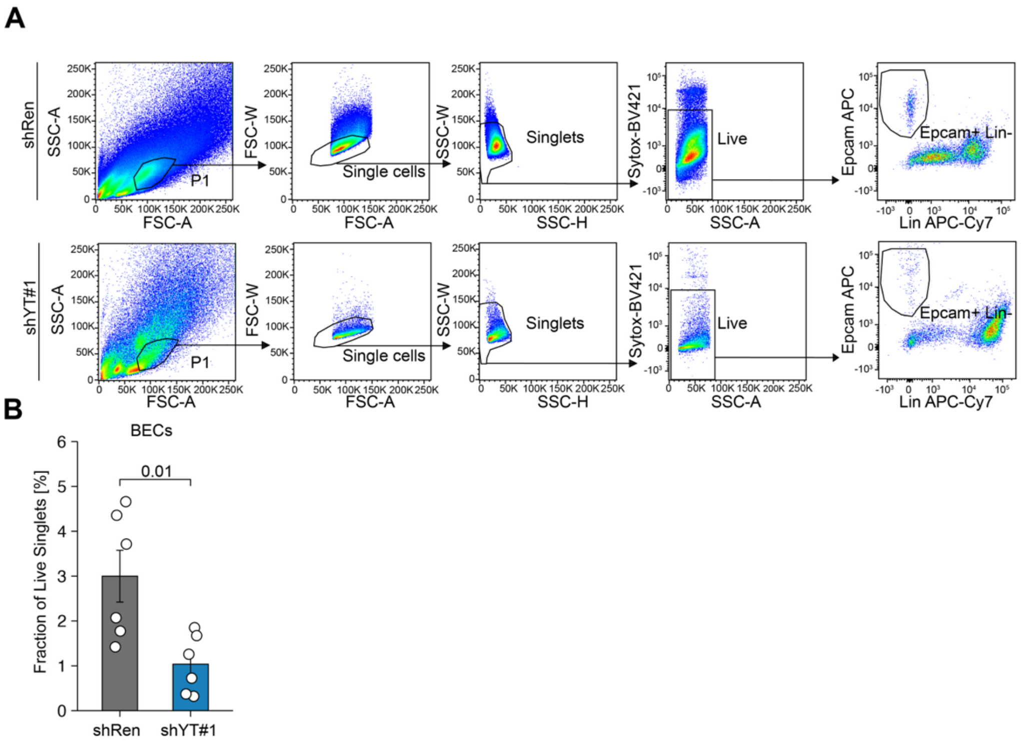


**Fig. S2. YAP/TAZ depletion reduces biliary epithelial cells (BECs) in the liver, as confirmed by fluorescence-activated cell sorting (FACS).**

(**A**, **B**) Representative FACS profiles and gating strategy from livers of shRen (top) and shYT#1 (bottom) mice. Quantification of BECs from live singlet populations (n = 6 per group).

Data represent mean ± SEM. Statistical analysis: Welch's t test with Benjamini-Hochberg correction.

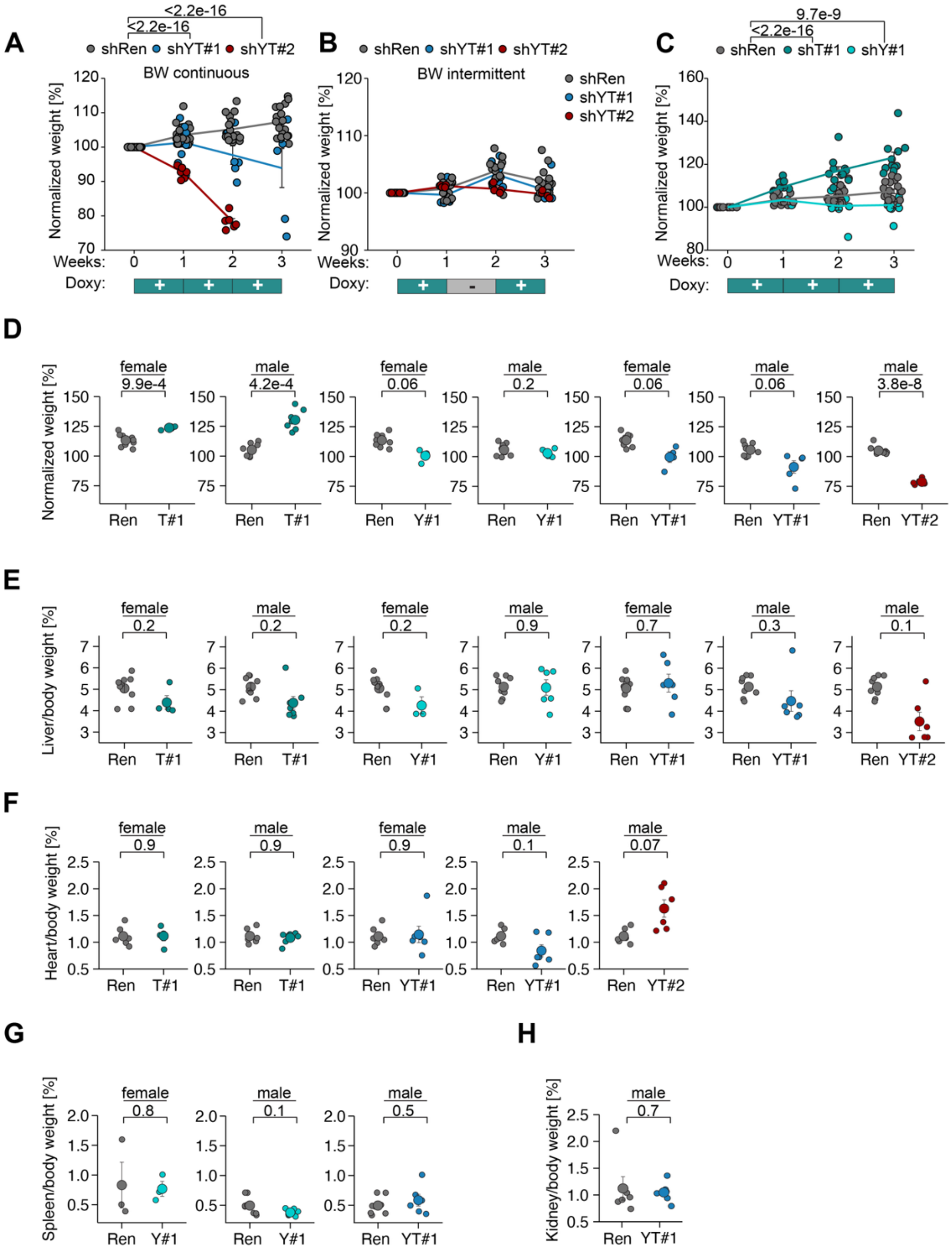


**Fig. S3. Body weight loss following constitutive YAP/TAZ depletion.**

(**A**) Change in body weight following continuous doxycycline-induced YAP/TAZ knockdown over 3 weeks (shRen, n = 17; shYT#1, n = 6; shYT#2, n = 6).

(**B**) Change in body weight following intermittent doxycycline administration (shRen, n = 15; shYT#1, n = 7; shYT#2, n = 4).

(**C**) Change in body weight over 3 weeks in mice with selective knockdown of YAP (shY#1), TAZ (shT#1), or both (shYT#1) (shRen, n = 15; shY#1, n = 9; shT#1, n = 11).

(**D**) Body weight normalized to baseline after 4 weeks of doxycycline treatment for shYT#1, shY#1, and shT#1, or 2 weeks for shYT#2, stratified by sex. Each dot represents one animal.

(**E**) Liver weight normalized to body weight after 4 weeks (shYT#1, shY#1, shT#1) or 2 weeks (shYT#2) of knockdown, stratified by sex.
(**F**) Heart weight normalized to body weight after 4 weeks (shYT#1, shT#1) or 2 weeks (shYT#2), stratified by sex.
(**G**) Spleen weight normalized to body weight after 4 weeks (shYT#1, shY#1) or 2 weeks (shYT#2), stratified by sex.
(**H**) Kidney weight normalized to body weight after 4 weeks of YAP/TAZ depletion (shYT#1), stratified by sex.
Data represent mean ± SEM. Statistical analysis: two-way ANOVA (A–C); Welch's t test with Benjamini-Hochberg correction (D–I).

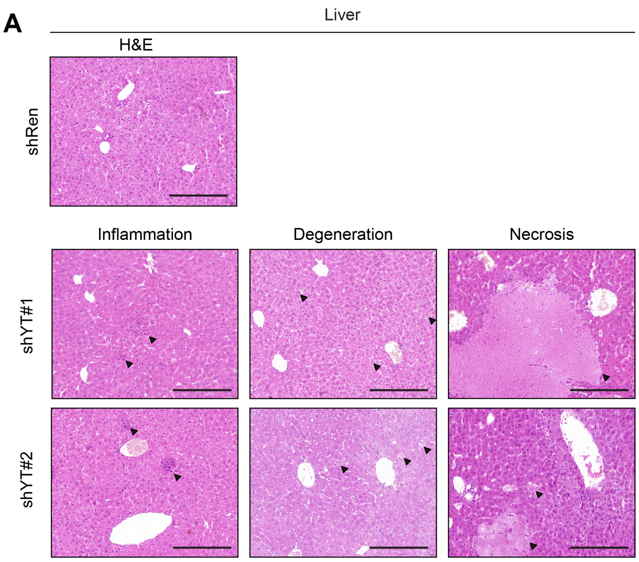


| | Inflammation | Degeneration | Necrosis |
|---|---|---|---|
| shRen | | | |
| shYT#1 | 4 out of 5 | 5 out of 5 | 2 out of 5 |
| shYT#2 | 2 out of 6 | 6 out of 6 | 6 out of 6 |

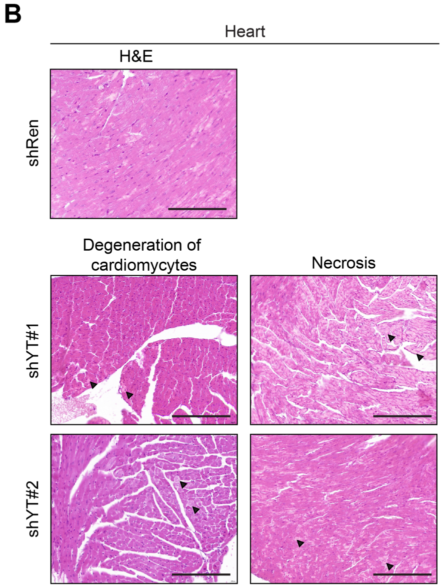


| | Degeneration | Necrosis |
|---|---|---|
| shRen | | |
| shYT#1 | 5 out of 5 | 5 out of 5 |
| shYT#2 | 5 out of 6 | 5 out of 6 |

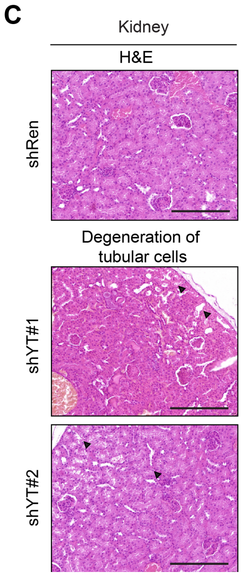


| | Degeneration |
|---|---|
| shRen | |
| shYT#1 | 5 out of 5 |
| shYT#2 | 6 out of 6 |

**Fig. S4. Histopathological manifestations in liver, heart, and kidney following long-term YAP/TAZ depletion.**

(**A**–**C**) Representative hematoxylin and eosin (H&E) staining of liver (A), heart (B), and kidney (C) sections from control (shRen) and YAP/TAZ-depleted (shYT#1, shYT#2) mice analyzed at their respective humane endpoints.

Liver sections show immune cell infiltration, parenchymal degeneration, and necrosis (arrows).

Heart tissue displays cardiomyocyte degeneration and necrotic areas (arrows).

Kidney sections reveal degeneration of tubular epithelial cells (arrows).

These findings reflect multi-organ pathology associated with sustained YAP/TAZ loss.

Scale bars, 200 μm.

Histological analysis revealed that shYT-deprived animals showed multifocal lesions in multiple organs.

<u>Cases shYT1#:</u>

- Liver showed severe multifocal to coalescent necrosis in 2 out of 5 cases; mild to moderate inflammation in 4 out of 5, composed of a multifocal lymphoplasmacytic infiltrate; mild to moderate hepatocyte degeneration with optically empty cytoplasm containing small, rounded vacuoles or granular material, which sometimes delocalizes the nucleus in the periphery in all cases.
- Heart showed mild to moderate compensatory hypertrophy of cardiomyocytes (degeneration) with variability in myofiber diameter, reduction in myofiber size (atrophy) with angular profile, and severe multifocal to coalescent necrosis in all cases.
- Kidney showed in all cases mild to moderate degeneration of tubular epithelial cells.

<u>Cases shYT2#:</u>

- Liver showed mild to moderate, multifocal to coalescent, necrosis in all cases; mild to moderate inflammation in 2 out of 6, composed of a multifocal lymphoplasmacytic infiltrate; mild to moderate hepatocyte degeneration with optically empty cytoplasm containing small, rounded vacuoles or granular material, which sometimes delocalizes the nucleus in the periphery in all cases.
- Heart showed mild to moderate compensatory hypertrophy of cardiomyocytes (degeneration) with variability in myofiber diameter, reduction in myofiber size (atrophy) with angular profile, and mild to moderate, multifocal to coalescent, necrosis in 5 out of 6 cases.
- Kidney showed in all cases mild to moderate degeneration of tubular epithelial cells.

A

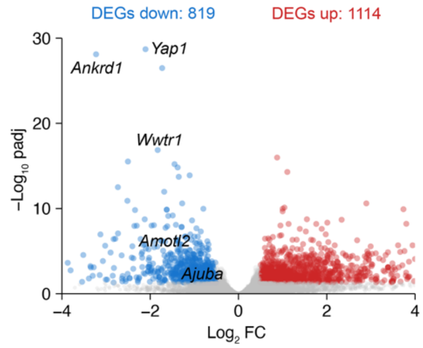


**Fig. S5. Transcriptomic analysis of acute YAP/TAZ depletion in CCA.**
(**A**) Acute YAP/TAZ depletion was induced in advanced CCA by doxycycline administration from day 25 to 33. Tumor cells were isolated from tumor-bearing livers by EpCAM-based enrichment and subjected to RNA-seq. Volcano plot shows downregulation of *Yap1*, *Wwtr1* (Taz), and canonical YAP/TAZ target genes (*Ankrd1*, *Amotl2*, *Ajuba*).

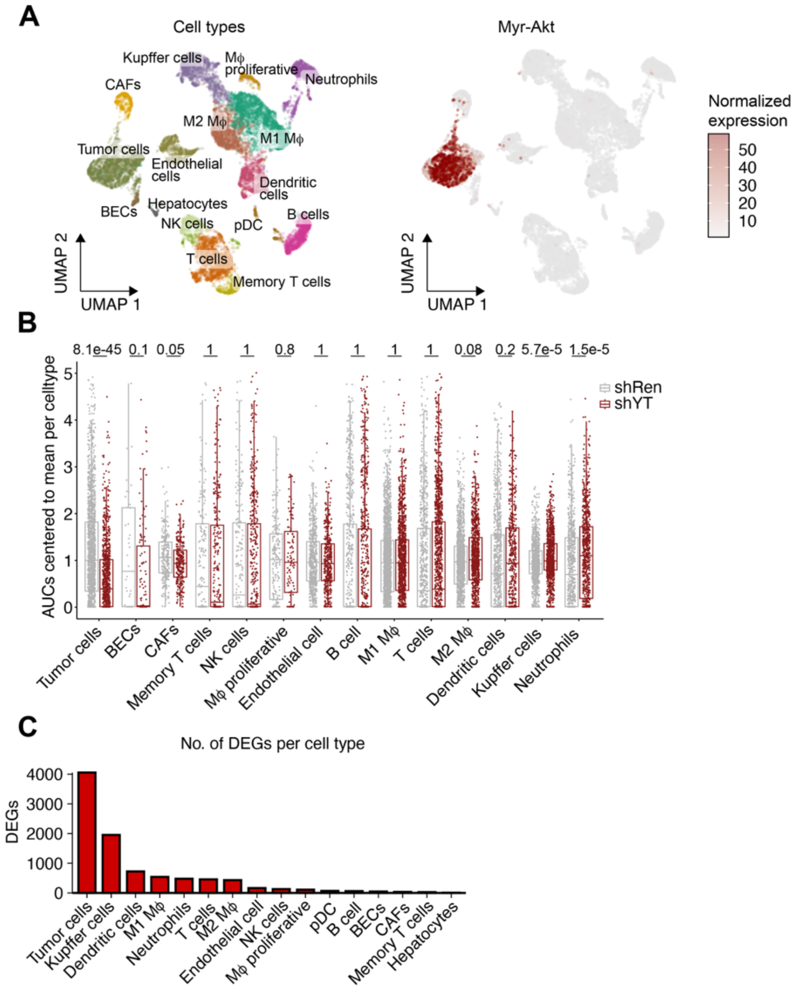


**Fig. S6. Tumor cells exhibit the strongest YAP/TAZ-dependent transcriptomic changes within the CCA tumor microenvironment (TME).**
(**A**) Acute YAP/TAZ depletion was induced in advanced CCA by doxycycline treatment from day 25 to 33. Single-cell RNA-seq was performed to assess transcriptomic changes within the TME. UMAP plots show an overview of identified cell types (left) and expression of Myr-AKT (right) to define the tumor cell cluster (shRen, n = 4; shYT#1, n = 2; shYT#2, n = 2).
(**B**) Box plots showing AUC scores for a conserved YAP transcriptional signature (18) in individual cell types from shRen and shYT samples. AUC values are centered to the mean of every celltype. Statistical analysis: Wilcoxon rank test
(**C**) Bar plot displaying the number of differentially expressed genes across cell types following YAP/TAZ depletion.

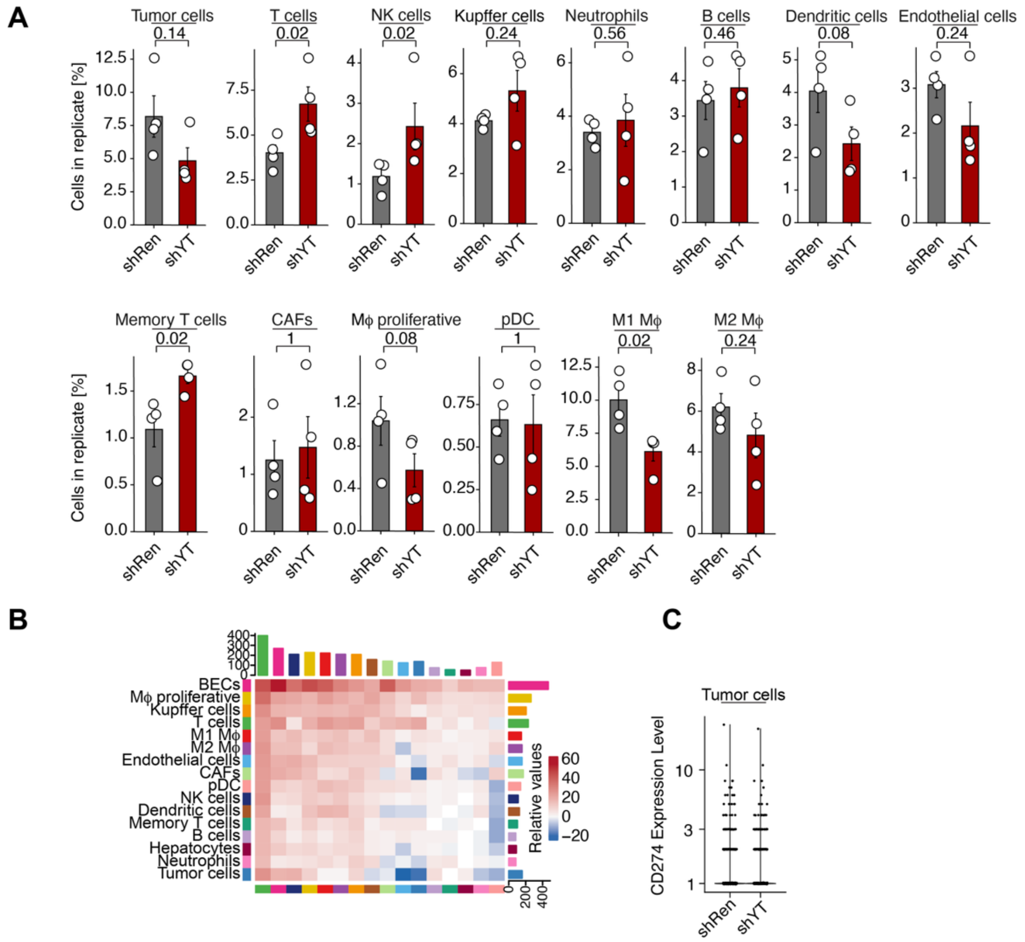


**Fig. S7. Quantitative changes in the tumor microenvironment (TME) of CCA following acute YAP/TAZ depletion.**

(**A**) Bar plots showing the proportion of each cell type across individual replicates from the scRNA-seq dataset comparing shRen (n = 4) with shYT (shYT#1, n = 2; shYT#2, n = 2). Statistical analysis: Welch's t test with Benjamini-Hochberg correction.

(**B**) Heatmap displaying the number of cell–cell interactions identified by CellChat analysis, stratified by cell type. Bar plots on the top and right indicate the total number of interactions sent and received, respectively, for each cell type.

(**C**) Violin plot illustrating the expression levels of *Cd274* (PD-L1) within the tumor cell cluster.

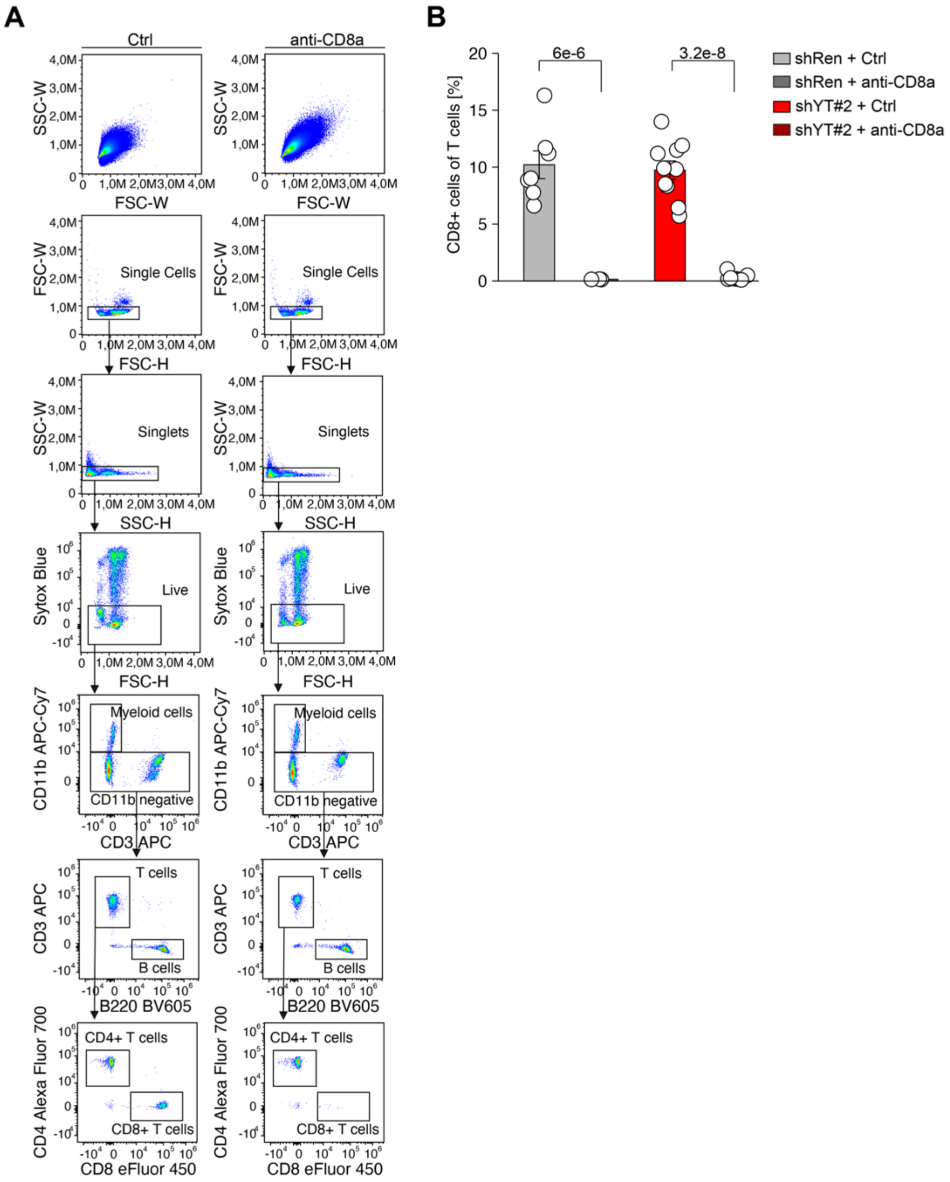


**Fig. S8. Validation of CD8 T cell depletion.**

(**A**, **B**) N-AKT–driven cholangiocarcinoma was induced via hydrodynamic tail vein injection (HDTVI), and CD8 T cells were depleted by intraperitoneal injection of monoclonal anti-CD8 antibodies on days 22 and 26 post-induction. Depletion efficiency was assessed on day 27. Representative FACS profiles and gating strategy are shown for peripheral blood from control (left) and anti-CD8–treated (right) mice. Quantification of $CD8^+$ T cells among live T cell populations (shRen Ctrl, n = 7; shRen anti-CD8, n = 3; shYT#2 Ctrl, n = 10; shYT#2 anti-CD8, n = 8).

Data represent mean ± SEM. Statistical analysis: one-way ANOVA with Tukey HSD post hoc test.

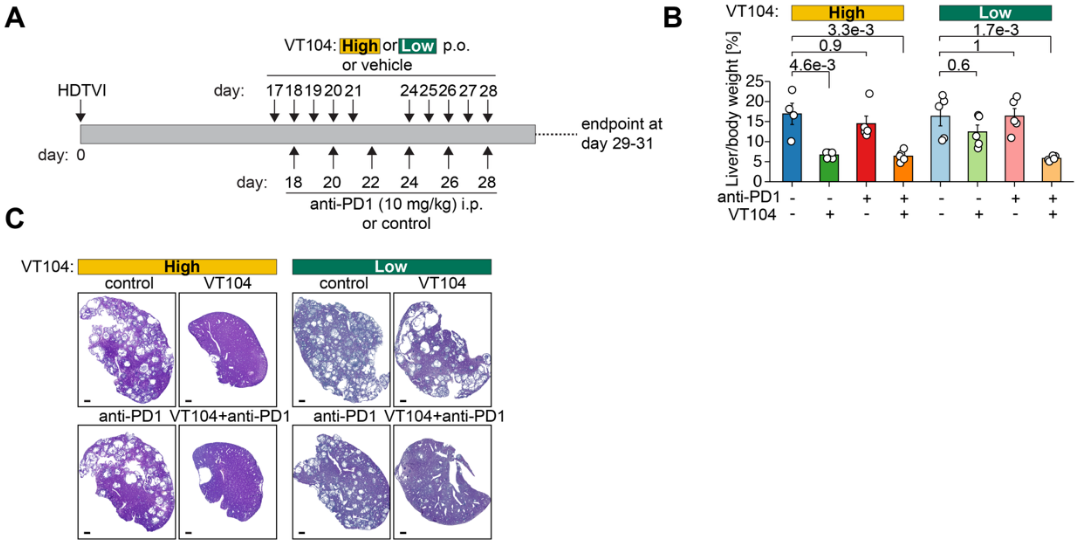


**Fig. S9. Low-dose TEAD inhibitor VT104 synergizes with anti–PD-1 therapy to reduce tumor burden in CCA.**

(**A**) Experimental schematic: N-AKT–driven cholangiocarcinoma was treated with two cycles of VT104 (administered from days 17–21 and 24–28 post-induction) at high (10 mg/kg) or low (1 mg/kg) doses, in combination with anti–PD-1 antibody (10 mg/kg, intraperitoneal). Tumor burden was assessed on days 29–31.

(**B**) Liver-to-body weight ratios for each treatment group (n = 5 per group; Ctrl of high-dose VT104, n = 4).

(**C**) Representative H&E-stained liver sections from the indicated treatment conditions. Scale bars, 1 mm.

Data represent mean ± SEM. Statistical analysis: one-way ANOVA with Tukey HSD post hoc test.

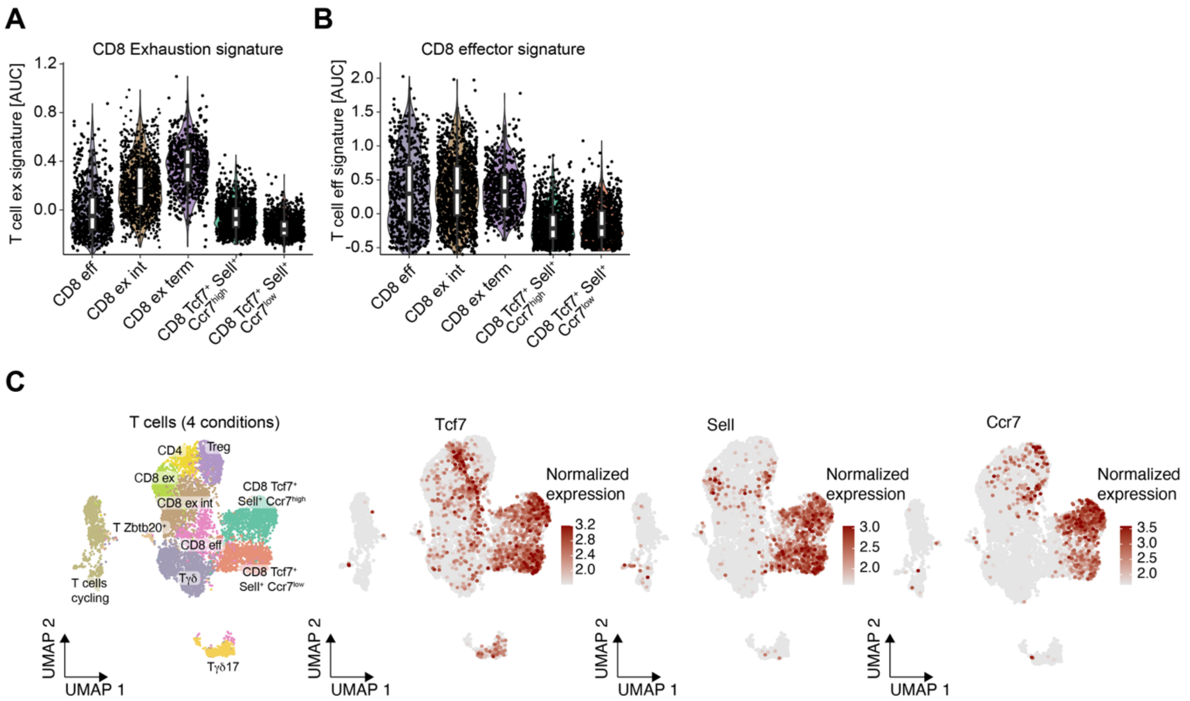


**Fig. S10. Annotation of T cell clusters based on marker gene expression.**
(**A**–**C**) Single-cell RNA-seq was performed following acute YAP/TAZ depletion in combination with anti–PD-1 therapy (see Fig. 4). T cell clusters were annotated based on expression of gene signatures associated with exhaustion (A), effector function (B), or naïve/memory-like markers (Tcf7, Sell, Ccr7) (C).

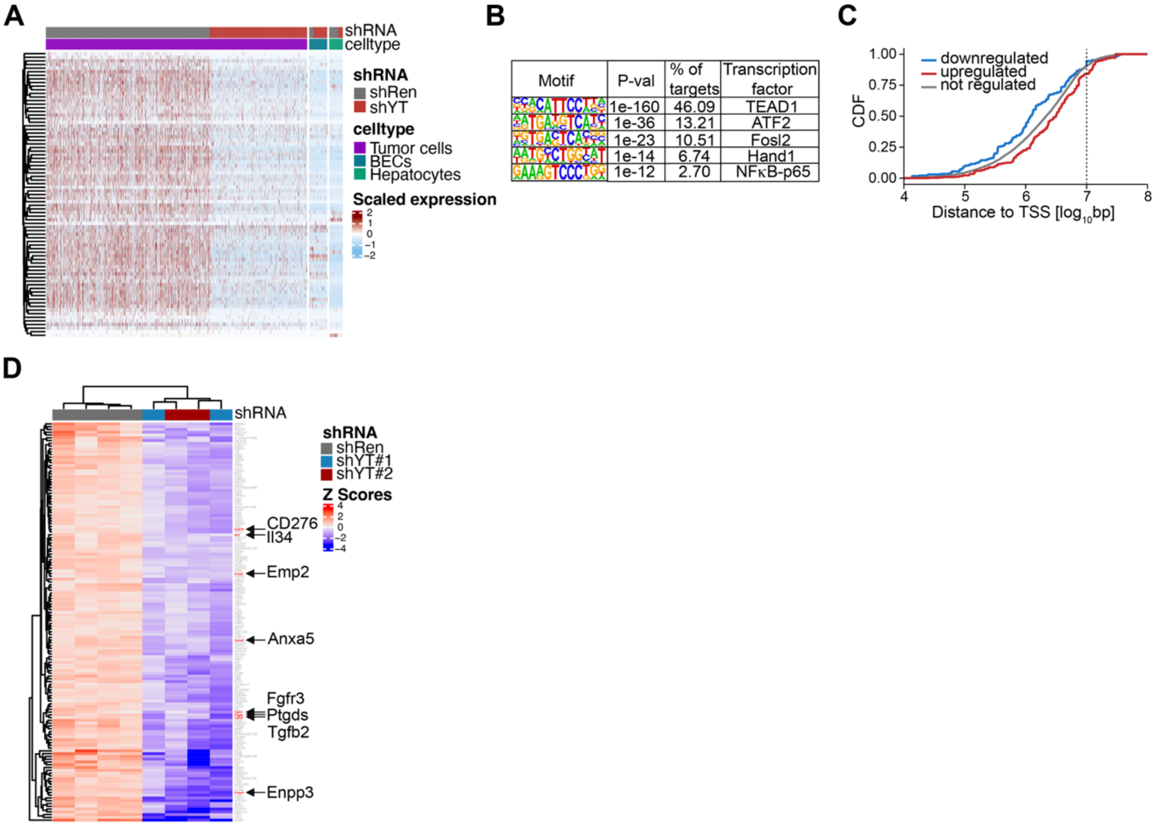


**Fig. S11. YAP/TAZ regulate a transcriptional program that maintains oncogenic features of CCA tumor cells.**

(**A**) Heatmap of YAP/TAZ-dependent transcriptional changes in tumor cells, showing no effects on gene expression in normal biliary epithelial cells (BECs) and hepatocytes. Genes upregulated in control tumor cells and downregulated in shYAP/TAZ-depleted cells are not differentially expressed in BECs or hepatocytes.

(**B**) HOMER motif analysis of TEAD1 CUT&RUN peaks, indicating enriched motifs, associated p-values, percentage of total peaks, and predicted transcription factors.

(**C**) Cumulative distribution function of TEAD1 CUT&RUN peaks plotted by distance to the nearest transcription start site (TSS), stratified by gene regulation status in RNA-seq (see Fig. S4).

(**D**) Heatmap showing the expression of 144 YAP/TAZ-regulated candidate genes—identified by combined CUT&RUN and RNA-seq analyses—in purified cholangiocarcinoma cells following YAP/TAZ knockdown (shRen, n = 4; shYT#1, n = 2; shYT#2, n = 2).

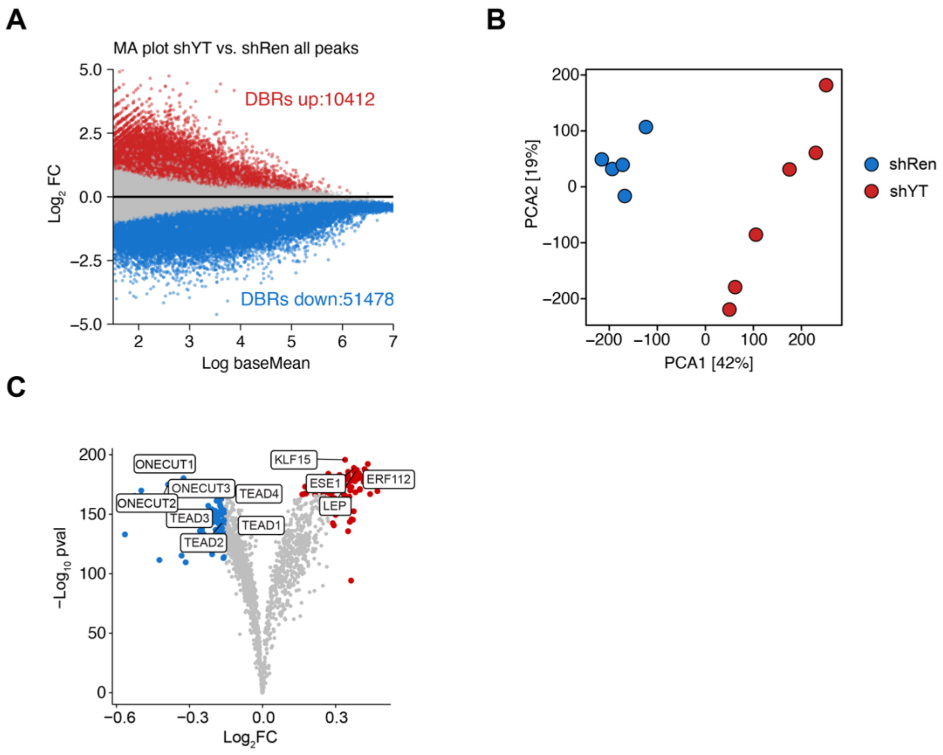


**Fig. S12. ATAC-seq reveals chromatin accessibility changes following acute YAP/TAZ depletion in CCA tumor cells.**

(**A**) MA plot showing differential chromatin accessibility in tumor cells after acute YAP/TAZ depletion, with 10,412 opening regions and 51,478 closing regions genome wide. Differential binding region, DBR.

(**B**) Principal component analysis (PCA) plot demonstrating separation between control (shRen) and YAP/TAZ-depleted (shYT#2) tumor cells based on ATAC-seq profiles.

(**C**) TOBIAS footprinting analysis of differentially accessible regions. Volcano plot displays significantly enriched transcription factor motifs in closing (blue) and opening (red) regions. (shRen, n = 5; shYT#2, n = 6).

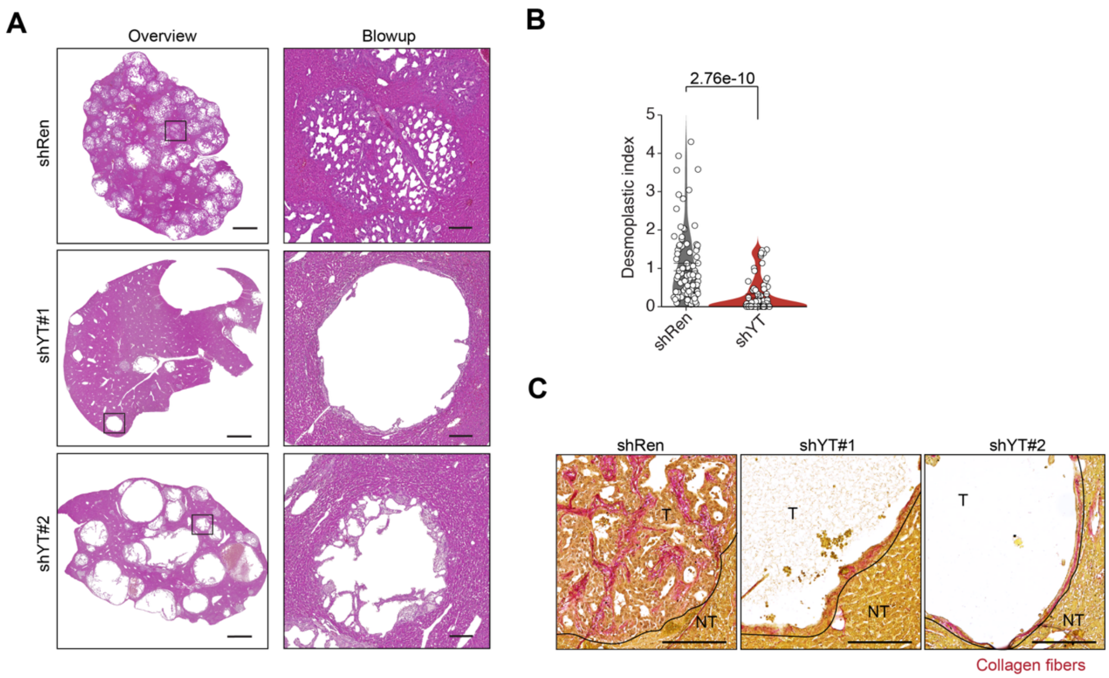


**Fig. S13. YAP/TAZ maintain the desmoplastic architecture of cholangiocarcinomas (CCAs).**

(**A**) Representative H&E-stained liver sections from tumor-bearing mice with acute YAP/TAZ depletion, showing low-magnification overviews (left) and higher-magnification zoom-ins (right). Scale bars, 2 mm (overview), 200 µm (zoom-in).

(**B**) Quantification of desmoplastic score in livers from control (shRen, n = 6) and YAP/TAZ-depleted animals (shYT#1, n = 2; shYT#2, n = 5). Statistical analysis: one-way ANOVA with Tukey HSD post hoc test.

(**C**) Representative images of Picro Sirius Red staining indicating collagen (red) in liver sections from tumor-bearing mice with acute YAP/TAZ depletion. Scale bars, 100 µm. NT, non-tumor tissue; T, tumor tissue.

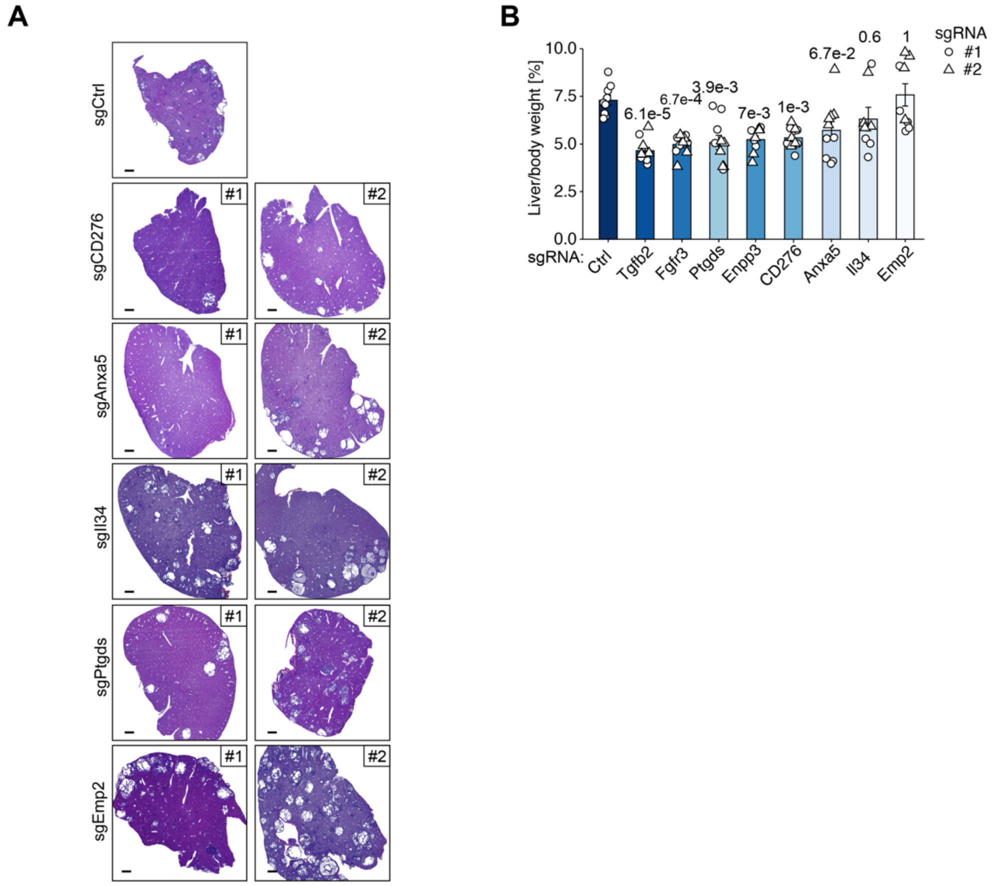


**Fig. S14. Functional validation of bona fide YAP/TAZ target genes in CCA growth.**
(**A**) Representative H&E-stained liver sections from tumor-bearing mice transduced with the indicated sgRNAs. Scale bars, 1 mm.
(**B**) Quantification of liver-to-body weight ratios in mice expressing the indicated sgRNAs (n = 5 per group; n = 4 for sgIl34#1 and #2; n = 9 for sgCtrl).
Statistical analysis: one-way ANOVA with Tukey HSD post hoc test.

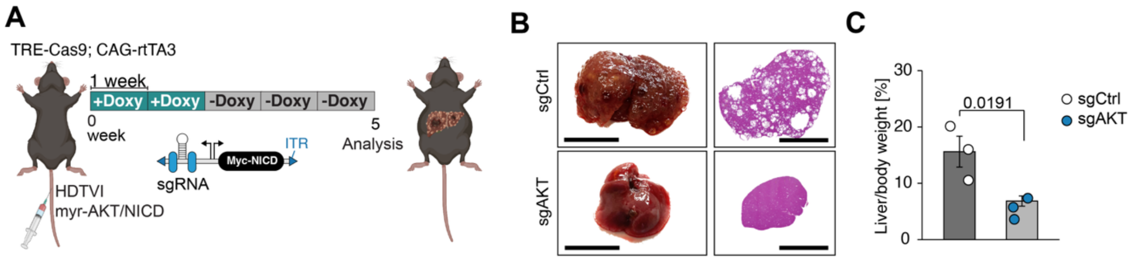


**Fig. S15. Knockout of the driver oncogene Myr-AKT inhibits N-AKT–induced CCA development.**

(**A**) Schematic of the inducible Cas9 mouse model used in combination with hydrodynamic tail vein injection (HDTVI) of sgRNA-expressing plasmids to assess the functional relevance of candidate YAP/TAZ targets in N-AKT–driven cholangiocarcinoma (CCA).

(**B**, **C**) Representative liver images (left) and H&E-stained liver sections (right) from mice with the indicated gene knockouts. Quantification of tumor area is shown. Scale bars, 1 cm (images), 500 µm (H&E). (n = 3 per group).

Statistical analysis: Welch's t test with Benjamini-Hochberg correction.

**Table S1. Transcriptional changes following acute depletion of YAP/TAZ in purified cholangiocarcinoma (CCAs) cells.**

This table lists differentially expressed genes after acute YAP/TAZ depletion in EpCAM enriched tumor cells. For each gene, the table provides the Entrez ID, Ensembl ID, gene symbol, base mean expression, log2 fold change, p-value, and adjusted p-value (padj).

**Table S2. Upregulated genes following acute depletion of YAP/TAZ in T cells.**

This table presents genes that are significantly upregulated in T cells upon acute YAP/TAZ depletion, as identified by single-cell RNA sequencing (scRNA-seq) analysis shown in Fig. 3C. These genes were subsequently used to stratify TCGA cholangiocarcinoma (CCA) patients into high- and low-expression groups, as shown in Fig. 3L. For each gene, the table includes the gene symbol, p-value, average log2 fold change, adjusted p-value (padj), and the corresponding Seurat cluster.

**Table S3. T cell exhaustion gene signature.**

This table lists the genes used to define the T cell exhaustion signature analyzed in Fig. 4E.

**Table S4. T cell effector gene signature.**

This table lists the genes used to define the T cell effector signature applied in the classification of T cells shown in Fig. 4M and Fig. 4N.

**Table S5. YAP/TAZ-driven transcriptional program sustaining oncogenic features of CCA.**

This table lists genes that are specifically downregulated in cholangiocarcinoma (CCA) tumor cells following YAP/TAZ depletion but not affected in non-malignant biliary epithelial cells (BECs) or hepatocytes. These data correspond to the heatmap shown in Fig. S10A.

**Table S6. Bona fide YAP/TAZ target genes identified in CCA.**

This table lists high-confidence YAP/TAZ target genes in cholangiocarcinoma (CCA), defined by the presence of a TEAD1 binding peak within a $10^7$ bp window near the transcription start site (TSS) (CUT&RUN analysis) and significant downregulation upon YAP/TAZ depletion (log2 fold change $< -1$; adjusted p-value $< 1\times10^{-3}$, bulk-RNAseq).

**Table S7. List of Antibodies**

| Name | Source | Identifier | Application, concentration |
|---|---|---|---|
| Anti-mouse PD-1 (cloneRMP1-14) | Bio X Cell | BE0146 | i.p. injection 10 mg/kg |
| Anti-mouse CD8 (clone YTS169) | Self produced | | i.p. 500 µg per mouse |
| SOX9 | Millipore | 5535 | IHC, 1:200 |
| CK19 | DSHB | TROMA-III | IHC, 1:15 |
| CD3 | Abcam | ab16669 | IHC, 1:150 |
| CD8 | Cell Signaling | 98941S | IHC, 1:200 |
| anti-EpCAM-APC | eBioscience | 17-5791-82 | FACS, 1:100 |
| anti-Mouse TER-119/Erythroid Cells-APC-Cy7 | BD Bioscience | 560509 | FACS, 1:100 |
| anti-CD11b-APC-Cy7 | BD Bioscience | 557657 | FACS, 1:100 |
| anti-Mouse CD45-APC-Cy7 | BD Bioscience | 557659 | FACS, 1:100 |
| H3K27Ac | Abcam | Ab4729 | CUT&RUN, 1:100 |
| H3K4me1 | Cell Signaling | 5326 | CUT&RUN, 1:100 |
| TEAD1 | Cell Signaling | 12292 | CUT&RUN, 1:100 |
| VINCULIN | Sigma-Aldrich | V9131 | WB, 1:10 000 |
| YAP (D8H1X) | Cell Signaling | 14074 | WB, 1:1000<br>IHC, 1:200<br>CUT&RUN, 1:100 |
| Goat Anti-Rabbit Immunoglobulins/HRP | Agilent | P044801-2 | WB, 1:5000<br>IHC, 1:1000 |
| Goat Anti-Mouse Immunoglobulins/HRP | Agilent | P044701-2 | WB, 1:5000<br>IHC, 1:1000 |
| TotalSeq-B0301 (Hashtag 1) | Biolegend | 155831 | scRNAseq, 1µg |
| TotalSeq-B0302 (Hashtag 2) | Biolegend | 155833 | scRNAseq, 1µg |
| TotalSeq-B0303 (Hashtag 3) | Biolegend | 155835 | scRNAseq, 1µg |
| TotalSeq-B0304 (Hashtag 4) | Biolegend | 155837 | scRNAseq, 1µg |

**Table S8. List of 70mer oligos for sgRNA cloning**

This table contains the sequences of 70-nucleotide (70mer) single-stranded DNA oligonucleotides used for the cloning of single guide RNAs (sgRNAs) into the psBbi-Myc-NICD-sgRNA vector. Each oligo includes necessary flanking sequences for compatibility with Gibson Assembly via SapI cutting site.

| Gene | Sequence 5’-3` | |
|---|---|---|
| | 70-mer | sgRNA target sequence |
| sgCtrl | ATCTTGTGGAAAGGACGAAACACCGATGTTG CAGTTCGGCTCGATGTTTTAGAGCTAGAAAT AGCAAGTT | ATGTTGCAGTTCGGCTCGAT |
| sgCD276#1 | ATCTTGTGGAAAGGACGAAACACCGCGCGT CCGAGTAACCGACGAGTTTTAGAGCTAGAAA TAGCAAGTT | CGCGTCCGAGTAACCGACGA |
| sgCD276#2 | ATCTTGTGGAAAGGACGAAACACCGGCGCG TCCGAGTAACCGACGGTTTTAGAGCTAGAAA TAGCAAGTT | GCGCGTCCGAGTAACCGACG |
| sgEmp2#1 | ATCTTGTGGAAAGGACGAAACACCGGCCGG TCAGCTCATTGATCTGTTTTAGAGCTAGAAA TAGCAAGTT | GCCGGTCAGCTCATTGATCT |

| | | |
|---|---|---|
| sgEmp2#2 | ATCTTGTGGAAAGGACGAAACACCGTTGGC GCCGGTCTGTATAGAGTTTTAGAGCTAGAAA TAGCAAGTT | TTGGCGCCGGTCTGTATAGA |
| sgEnpp3#1 | ATCTTGTGGAAAGGACGAAACACCGATATTA CGAGTCCTGATTCGGTTTTAGAGCTAGAAAT AGCAAGTT | ATATTACGAGTCCTGATTCG |
| sgEnpp3#2 | ATCTTGTGGAAAGGACGAAACACCGACGTG CAATTTATTCCGTTGGTTTTAGAGCTAGAAAT AGCAAGTT | ACGTGCAATTTATTCCGTTG |
| sgFgfr3#1 | ATCTTGTGGAAAGGACGAAACACCGAGCCG GGCAATCCGGACAAGGTTTTAGAGCTAGAA ATAGCAAGTT | AGCCGGGCAATCCGGACAAG |
| sgFgfr3#2 | ATCTTGTGGAAAGGACGAAACACCGGGTAT AGTTGCCACGATCGGGTTTTAGAGCTAGAAA TAGCAAGTT | GGTATAGTTGCCACGATCGG |
| sgIl34#1 | ATCTTGTGGAAAGGACGAAACACCGGACCT TACAGGCTACCTTCGGTTTTAGAGCTAGAAA TAGCAAGTT | GACCTTACAGGCTACCTTCG |
| sgIl34#2 | ATCTTGTGGAAAGGACGAAACACCGTCTTG GGATCCTACTTGACGGTTTTAGAGCTAGAAA TAGCAAGTT | TCTTGGGATCCTACTTGACG |
| sgPtgds#1 | ATCTTGTGGAAAGGACGAAACACCGATTGA GGCCGCCTTCTGTGGGTTTTAGAGCTAGAA ATAGCAAGTT | ATTGAGGCCGCCTTCTGTGG |
| sgPtgds#2 | ATCTTGTGGAAAGGACGAAACACCGCAGAG CGTACTCGTCATAGTGTTTTAGAGCTAGAAA TAGCAAGTT | CAGAGCGTACTCGTCATAGT |
| sgAnxa5#1 | ATCTTGTGGAAAGGACGAAACACCGCATTG CTTCGGGATGTCAACGTTTTAGAGCTAGAAA TAGCAAGTT | CATTGCTTCGGGATGTCAAC |
| sgAnxa5#2 | ATCTTGTGGAAAGGACGAAACACCGTAGGC ATCGTAGAGTCGTGAGTTTTAGAGCTAGAAA TAGCAAGTT | TAGGCATCGTAGAGTCGTGA |
| sgTgfb2#1 | ATCTTGTGGAAAGGACGAAACACCGCGAGG AGTACTACGCCAAGGGTTTTAGAGCTAGAAA TAGCAAGTT | CGAGGAGTACTACGCCAAGG |
| sgTgfb2#2 | ATCTTGTGGAAAGGACGAAACACCGTGGAT CAGTTTATGCGCAAGGTTTTAGAGCTAGAAA TAGCAAGTT | TGGATCAGTTTATGCGCAAG |

### Table S9. Oligos used for ATACseq library amplification

| Primer | Sequence 5’-3` | |
|---|---|---|
| | sequence | barcode |
| Ad1.1 | AATGATACGGCGACCACCGAGATCTACACTAGATCGCTCGTCGGCA GCGTCAGATGTGTAT | TAGATCGC |
| Ad1.2 | AATGATACGGCGACCACCGAGATCTACACCTCTCTATTCGTCGGCAG CGTCAGATGTGTAT | CTCTCTAT |
| Ad1.3 | AATGATACGGCGACCACCGAGATCTACACTATCCTCTTCGTCGGCAG CGTCAGATGTGTAT | TATCCTCT |
| Ad1.4 | AATGATACGGCGACCACCGAGATCTACACAGAGTAGATCGTCGGCA GCGTCAGATGTGTAT | AGAGTAGA |
| Ad1.5 | AATGATACGGCGACCACCGAGATCTACACGTAAGGAGTCGTCGGCA GCGTCAGATGTGTAT | GTAAGGAG |

| | | |
|---|---|---|
| Ad1.6 | AATGATACGGCGACCACCGAGATCTACACACTGCATATCGTCGGCAG CGTCAGATGTGTAT | ACTGCATA |
| Ad2.1 | CAAGCAGAAGACGGCATACGAGATTCGCCTTAGTCTCGTGGGCTCG GAGATGTG | TAAGGCGA |
| Ad2.2 | CAAGCAGAAGACGGCATACGAGATCTAGTACGGTCTCGTGGGCTCG GAGATGTG | CGTACTAG |

### Table S10. Oligos used for qRT-PCR

| **Primer** | **Sequence 5'-3`** | |
|---|---|---|
| | Forward | Reverse |
| Yap1 | TGAGATCCCTGATGATGTACCAC | TGTTGTTGTCTGATCGTTGTGAT |
| Wwtr1 | GAAGGTGATGAATCAGCCTCTG | GTTCTGAGTCGGGTGGTTCTG |
| Ctgf | GGGCCTCTTCTGCGATTTC | ATCCAGGCAAGTGCATTGGTA |
| Cyr61 | CTGCGCTAAACAACTCAACGA | GCAGATCCCTTTCAGAGCGG |

### Table S11. Commercial assays

| **Name** | **Source** | **Identifier** |
|---|---|---|
| innuMix qPCR DSGreen Standard | Analytik Jena | 845-AS-1300200 |
| NEBNext® Ultra II RNA Library Prep kit for Illumina | NEB | E7770 |
| NEBNext® Poly(A) mRNA Magnetic Isolation Module | NEB | E7490 |
| NEBNext Multiplex Oligos for Illumina | NEB | E6440 |
| Chromium Next GEM Single Cell 3' Reagent Kits v3.1 | 10x Genomics | PN-1000268 |
| GEXSCOPE Single Cell RNA Library Kit V2 | Singleron | SD: 4180011/ 4180012 |
| NEBNext Ultra II DNA Library Prep Kit for Illumina | NEB | E7645 |
| Illumina Tagment DNA Enzyme and Buffer Small Kit | Illumina | 20034197 |
| DNA Clean & Concentrator-5 (Uncapped) | ZYMO Research | D4004 |
| NEBNext High-Fidelity 2X PCR Master Mix | NEB | M0541 |
| Click-iT™ EdU Cell Proliferation Kit | Thermo Fisher Scientific | C10337 |
| Red Blood Cell Lysis Solution | Miltenyi | 130-094-183 |
| Debris Removal Solution | Miltenyi | 130-109-398 |
| Dead Cell Removal Kit | Miltenyi | 130-090-101 |
| RNeasy Mini Kit | Quiagen | 74104 |
| RNase-Free DNase Set | Quiagen | 79254 |

### Table S12. Chemicals and enzymes

| **Name** | **Source** | **Identifier** |
|---|---|---|
| 0.25% Trypsin | Thermo Fisher Scientific | 25200056 |
| 37% Formaldehyde | Carl Roth | 4979.1 |
| AdDMEM/F-12 | Thermo Fisher Scientific | 12634010 |
| Antigen Retrieval Buffer 100x Citrate pH 6 | abcam | Ab64236 |
| Antigen Retrieval Buffer 100x Tris-EDTA, pH 9 | abcam | Ab93684 |
| BioMag Plus concanavalin A | Polysciences | 86057-10 |
| Bicine | Sigma-Aldrich | 215589 |
| Bis-Tris | Carl Roth | 9140.3 |
| Bovine Serum Albumin (BSA) | Carl Roth | 8076.1 |

| | | |
|---|---|---|
| CaCl2 | Carl Roth | CN92.1 |
| Chlorobutanol | Sigma-Aldrich | 112054 |
| Clarity Max Western ECL Substrate | Bio Rad | 1705062 |
| Collagenase P | Sigma Aldrich | 11213857001 |
| Digitonin | Sigma-Aldrich | D141 |
| Direct Red 80 (Sirius Red) | Sigma Aldrich | 365548 |
| DMEM (GlutaMAX™) | Thermo Fisher Scientific | 61965059 |
| doxycycline-containing chow (625 mg/kg doxycycline hyclate | Sniff | 326652 |
| DNaseI | Roche | 05952077103 |
| EDTA | Merck | 1.08452.1000 |
| EGTA | Carl Roth | 3054.3 |
| Eosin G-solution | Carl Roth | X883.2 |
| EpCAM-APC antibody | eBioscience | 17-5791-82 |
| Fc receptor blocking reagent, mouse | Miltenyi | 130-092-575 |
| Fetal Bovine Serum | Gibco | F7524-500ML |
| Glycerol | Carl Roth | 3783.2 |
| Glycine | Carl Roth | 3908.3 |
| GlycoBlue™ Coprecipitant | Invitrogen | AM9515 |
| Hematoxylin solution (Meyer) | Carl Roth | T865.1 |
| Hematoxylin solution A (Weigert) | Carl Roth | X906.1 |
| Hematoxylin solution B (Weigert) | Carl Roth | X907.1 |
| HEPES | Sigma-Aldrich | H3375 |
| Hoechst 33343 | Sigma-Aldrich | B2261 |
| Hydrogen peroxide solution (30%) | Honeywell | 216763 |
| Immobilon-P Transfer Membrane (0.45 µm) | Merck | IPVH00010 |
| Isopropanol | Carl Roth | T902.1 |
| KCl | Carl Roth | 6781.2 |
| M-MLV Reverse Transcriptase | Promega | M1705 |
| MgCl2 | Carl Roth | KK36.3 |
| Methanol | Carl Roth | 4627.2 |
| MnCl2 | Carl-Roth | 0276.1 |
| NP-40 | Sigma Aldrich | 74385 |
| Penicillin/Streptomycin | Thermo Fisher Scientific | 15140122 |
| peqGOLD TriFast™ | VWR | 30-2010 |
| Picric acid solution (1.3 % in H2O) | Sigma Aldrich | P6744 |
| Potassium dihydrogen phosphate (K2HPO4) | Carl Roth | P018.1 |
| Powdered Milk | Carl Roth | T145.2 |
| Pronase | Sigma Aldrich | 10165921001 |
| Protease Inhibitor Cocktail (PIC) | Sigma-Aldrich | P8340 |
| Random primer p(dN)6 | Sigma-Aldrich | 11034731001 |
| Ringer-Lactate solution | WDT | - |
| RNase A | Quiagen | 1032722 |
| ROTI®Phenol | Carl Roth | 0038.2 |
| SDS Pellets | Carl Roth | CN30.2 |
| Sodium bisulfite (NaHSO4) | Sigma-Aldrich | 243973 |
| Sodium chloride (NaCl) | Carl Roth | P029.3 |
| Sodium-deoxycholate | Sigma Aldrich | D6750 |
| Spermidine | Sigma-Aldrich | S0266 |
| Tris | Carl Roth | A411.2 |
| Tris-HCl | Sigma-Aldrich | T3253 |
| Tween-20 | Carl Roth | 9127.1 |

| | | |
|---|---|---|
| Vector® ImmPACT DAB Peroxidase (HRP)Sub. | Vector Laboratories | SK-4105 |

### Table S13. Plasmid constructs

| Name | Backbone | Insert |
|---|---|---|
| **Retrovirus constructs** | | |
| pRS-Puro sh p19 Arf | pRetro-Super-Puro | Sh p19Arf (mouse) |
| **HDTVI plasmids** | | |
| pCMV(CAT)T7-SB100 | pCMV | SB100 |
| pSBbi-w/oPuro-Myr-AKT-HA | pSBbi-w/oPuro | Myr-AKT-HA |
| pSBbi-w/oPuro-Myc-NICD | pSBbi-w/oPuro | Myc-NICD |
| psBbi-Myc-NICD-sgRNA | pSBbi-w/oPuro | Myc-NICD; sgRNA |
